# Valency-driven division of labor balances chromatin compaction and structural plasticity

**DOI:** 10.64898/2026.01.06.698048

**Authors:** Jiahu Tang, Xiakun Chu

## Abstract

Chromatin folding is regulated by multivalent protein complexes and condensates, yet how the multivalent binding quantitatively controls chromatin compaction, domain organization, and cell-to-cell heterogeneity remains unresolved. Here we develop the Chromatin-associated Protein Complex Maximum Entropy Model (CPC-MEM), a data-driven, physics-based polymer framework in which Hi-C contacts are realized through an explicit, finite pool of diffusing chromatin-associated protein complexes (CPCs) with prescribed CPC-chromatin interaction valency. When fitted to Hi-C, CPC-MEM generates chromatin structural ensembles that simultaneously reproduce population-averaged contact maps and single-cell super-resolution distance statistics. We uncover a clear division of labor across valency: low-valency CPCs act as abundant linkers that compact chromatin, homogenize folding, and refine local structure, whereas high-valency CPCs form sparse co-bridging hubs that nucleate chromatin domains and increase conformational heterogeneity. Mixtures of valencies naturally produce a “hub-and-matrix” chromatin architecture that optimally balances compaction with structural plasticity. Our study provides a quantitative mechanism by which locus-specific chromatin interaction preferences, together with the valency and abundance of CPCs and condensates in the nucleus, jointly shape functional 3D genome organization.

---

In cell nucleus, the genome is densely compacted yet nonrandomly folded to regulate which genes are activated and which are silenced [1, 2]. This organization is hierarchical, spanning kilobase-scale chromatin loops [3], megabase-scale topologically associating domains (TADs) [4], and chromosome-scale compartments A/B [5]. Such a complex architecture is not an intrinsic property of the chromatin polymer alone, but is actively regulated by a myriad of chromatin-associated protein complexes (CPCs) [6]. Examples range from structural maintenance of chromosomes (SMC) complexes that extrude chromatin loops [7, 8] to transcriptional regulators that can assemble the chromatin into mesoscale clusters or phase-separated condensates [9, 10]. The binding of CPC with chromatin converts local biochemical affinities into effective long-range couplings along the polymer, producing emergent outcomes such as bridging-induced attraction and selective clustering of regulatory elements, thereby reshaping chromatin accessibility and transcriptional output [11–14]. Despite these advances, how the CPC interaction collectively and quantitatively tunes genome folding remains poorly understood.

This knowledge gap is sharpened by an apparent tension between population-level reproducibility and single-cell variability in chromatin biology. Population-averaged experiments such as Hi-C reported stable chromatin architectural features across many cells [1]. In contrast, single-cell approaches and super-resolution imaging reveal that chromatin conformations are highly dynamic and heterogeneous, with stochastic fluctuations and substantial cell-to-cell variability [15, 16]. This discrepancy is highlighted by the long-noted “Hi-C-FISH paradox”, where contact frequencies and spatial distances cannot be reconciled without explicitly accounting for chromatin ensemble heterogeneity (Fig. S1) [17, 18]. Together, these observations have raised a fundamental biophysical question: how can a system governed by stochastic molecular interactions generate robust, cell-type-specific structural patterns, while retaining sufficient conformational plasticity for regulatory adaptation [19]? Addressing this question requires quantitative rules that connect molecular-scale binding interactions to emergent genome architecture and dynamics [20–23]. As many CPCs and condensate scaffolds present multiple binding modules [24–26], this enables them to impose constraints with distinct topologies on chromatin, ranging from predominantly pairwise bridging to multiplex co-bridging among several loci. In principle, changing the interaction valency can redistribute how a finite connectivity budget is converted into geometric constraints [27], potentially tuning the balance between chromatin compaction, domain formation, and cell-to-cell structural heterogeneity. However, a underlying understanding of how CPC valency encodes this balance is still lacking.

To quantitatively dissect the valency-dependent mechanisms described above, a framework is required to connect microscopic CPC binding to macroscopic, data-consistent structural ensembles. Unfortunately, most existing approaches in chromatin modeling emphasize one aspect at the expense of the other [28]. Top-down approaches based on the maximum-entropy principle can reconstruct structural ensembles consistent with Hi-C contact probabilities [29–31]. However, in conventional maximum-entropy models (MEMs), chromatin organization is typically encoded through effective, pairwise interactions, so the interaction valency and binding of the underlying CPCs are not represented explicitly. Conversely, bottom-up polymer models, such as the Strings-and-Binders-Switch (SBS) framework [32] and loop-extrusion models [7, 33], provide mechanistic insight into many-body bridging and the emergence of chromatin domains and loops. Yet, in most implementations these models treat CPC connectivity in an implicit manner, and are not designed to systematically infer the effective interaction valency of CPCs directly from experimental data [34]. As a result, it remains challenging to mechanistically relate experimentally measured chromatin structural characteristics to the microscopic interaction mediated by CPCs.

To meet this challenge, we develop a maximum-entropy polymer framework, termed CPC-MEM, that reproduces population Hi-C contact probabilities while explicitly representing CPCs and controlling their interaction valency. In both the conventional chromatin-chromatin maximum-entropy model (CC-MEM) and our CPC-mediated extension (CPC-MEM), the chromatin segment is represented as a coarse-grained semiflexible polymer consisting of *N* beads. The chromatin conformational ensemble is defined by the same maximum-entropy energy functional [29],

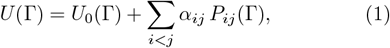

where *U*_0_(Γ) is a reference potential capturing baseline polymer physics (chain connectivity, bending rigidity, and excluded-volume interactions), {*α*_*ij*_} are Lagrange multipliers inferred from Hi-C data, and *P*_*ij*_(Γ) is the model-specific contact observable between loci *i* and *j*.

In CC-MEM, contacts are assumed to arise from direct proximity between chromatin beads (Fig. 1a). Accordingly,

**Figure 1.**
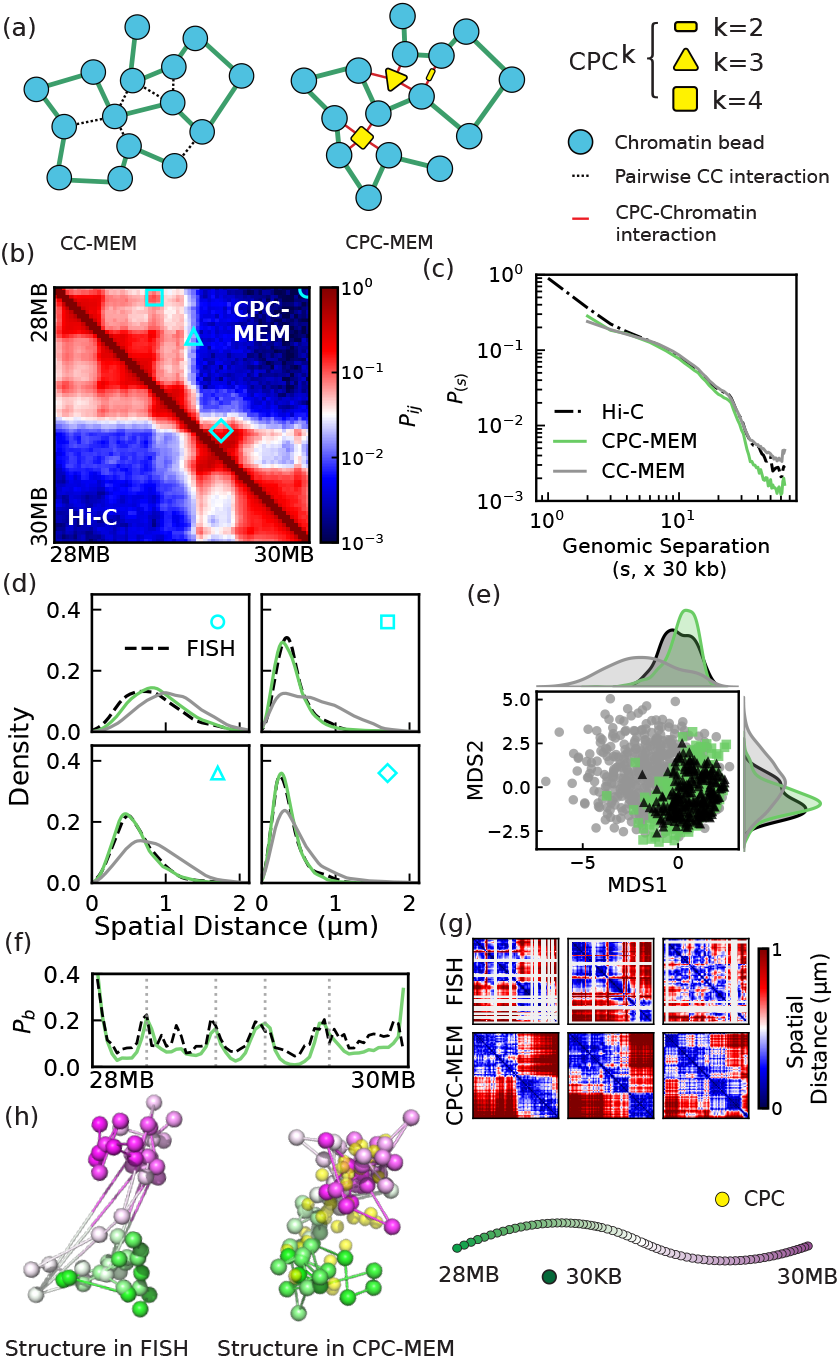
CPC-MEM framework and validation against population and single-cell measurements. (a) Schematic of CC-MEM and CPC-MEM. (b) Hi-C contact-probability map for the modeled chromatin region (Chr21: 28–30 Mb). Cyan markers indicate the four locus pairs used for the single-cell distance comparisons in (d). (c) Mean contact probability *P*(*s*) as a function of genomic separation *s*. (d) Single-cell spatial-distance distributions for the four locus pairs highlighted in (b). (e) Multidimensional scaling (MDS) embedding of conformations using pairwise-distance root-mean-square -deviation (dRMSD) as the dissimilarity metric. (f) Insulation boundary probability along the genomic coordinate. Grey dashed lines denote candidate TAD boundaries. (g) Representative single-cell spatial-distance matrices from experiment and from CPC-MEM. (h) Representative 3D conformations from experiment and from CPC-MEM, colored by genomic coordinate.

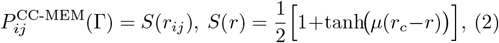

where 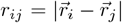, and *S*(*r*) switches from 1 to 0 controlled by a cutoff distance *r*_*c*_, with steepness set by *µ*. With this definition, *α*_*ij*_ reflects an effective, direct pairwise interaction between loci *i* and *j*.

In CPC-MEM, we introduce a finite pool of *M* explicit CPCs. Each CPC is represented as a spherical bead that freely diffuses and interacts with the chromatin polymer (Fig. 1a). The configuration now includes both chromatin and CPC coordinates, 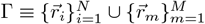. We define chromatin contacts through CPC-mediated bridging. The contribution of CPC *m* to bridging loci *i* and *j* is

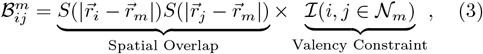

where the first two factors enforce simultaneous spatial proximity to CPC *m*. The indicator ℐ implements a strict valency constraint: ℐ = 1 only if both loci belong to 𝒩_*m*_, defined as the set of the *k* nearest chromatin beads to CPC *m* (so that each CPC can engage at most *k* loci at a time). The CPC-mediated contact observable *P*_*ij*_ is then defined as the probability that loci *i* and *j* are simultaneously bridged by at least one CPC in the current configuration.

By assuming statistical independence among the CPC binding to chromatin, this is formulated as

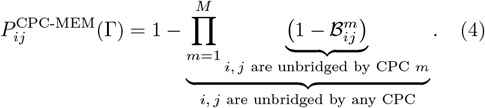

With this definition, the inferred parameters {*α*_*ij*_} in CPC-MEM acquire a different physical meaning than in CC-MEM. Rather than prescribing a direct chromatin-chromatin attraction, {*α*_*ij*_} quantifies the propensity for loci *i* and *j* to be bridged by a shared CPC under an explicit, valency-controlled binding topology. Given the resulting potential *U*(Γ), we generate the conformational ensemble of the system using Monte Carlo sampling (see details in Supplemental Material) [35].

We modeled a 2 Mb region on chromosome 21 (Chr21: 28– 30 Mb) in IMR90 cells at 30 kb resolution, corresponding to a semiflexible polymer of *N* = 65 chromatin beads. We focus on an intermediate-valency setting with *k* = 4 and *M* = 25 (denoted 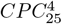) and compare with CC-MEM. Both models were trained to match the same Hi-C contact probabilities (Fig. S2). We further evaluated the resulting chromatin ensembles using independent single-cell spatial measurements from super-resolution chromatin tracing [16].

At the population level, CPC-MEM and CC-MEM both reproduce the Hi-C characteristics with high fidelity (Fig. 1b,c). However, the models diverge at the single-cell level. CPC-MEM accurately captures the traced spatial-distance distributions for the four locus pairs (Fig. 1d), whereas CC-MEM shows systematic deviations, indicating that fitting Hi-C alone does not fix the correct single-cell geometry. This difference is also evident in the dRMSD-based MDS embedding: the CC-MEM ensemble occupies a region largely disjoint from the experimentally sampled conformational space, while the CPC-MEM ensemble overlaps closely with the tracing-derived distribution (Fig. 1e). Beyond pairwise distances, CPC-MEM recapitulates stochastic domain organization across cells, reproducing the variability of TAD boundary locations quantified by the boundary probabilities *P*_*b*_ (Fig. 1f) and the cell-to-cell differences visible in representative distance matrices and 3D conformations (Fig. 1g,h). Together, these results show that introducing explicit, valency-controlled CPCs enables a MEM to match both population Hi-C and single-cell chromatin architecture statistics.

To isolate how CPC valency shapes the inferred chromatin ensemble, we varied the interaction valency *k* in CPC-MEM while keeping the same Hi-C map as the training target (Fig. S3). To ensure a fair comparison across different *k*, we fixed the total connectivity budget, 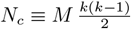, to a constant value (*N*_*c*_ ≃ 150), with adjusting *M* accordingly. Under this constant-*N*_*c*_ condition, the chromatin structure exhibits a clear valency-driven transition: low-valency systems (e.g., 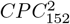) collapse chromatin into overly compact, homogeneous conformations, whereas high-valency systems (e.g., 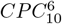) generate overly expanded, heterogeneous chromatin ensembles characterized. Overall, the intermediate-valency systems yield the closest agreement with the single-cell FISH benchmarks, indicating that realistic chromatin folding reflects a valency-tunable balance between compaction and heterogeneity. We find that these trends are robust to alternative constraint conditions. In particular, when CPC abundance is held fixed (a constant *M*) rather than *N*_*c*_, intermediate valency again provides the best match to the single-cell distance and heterogeneity statistics, while the low-valency limit becomes connectivity-limited and even cannot realize sufficient links to support the same Hi-C constraints (Fig. S4).

To further dissect valency-dependent roles, we performed *in silico* CPC-depletion simulations, mimicking the reduction of CPC concentrations in nucleus. Starting from the saturated reference state (*N*_*c*_ ≃ 150), we fixed the {*α*_*ij*_} and progressively reduced the number of realized CPC-mediated links and quantified the structural response (Fig. 2 and Fig. S5). Depletion degrades the reference ensemble for all *k*, but the failure mode depends strongly on valency. In the low-valency limit (*k* = 2), CPCs act as constraint-efficient pairwise bridges that percolate through the polymer. As a result, they provide the dominant contribution to overall compaction and to maintaining boundary insulation (Fig. 2a,b). Consistent with a percolating “matrix” of constraints, locus fluctuations remain uniformly suppressed along the segment (Fig. 2d).

**Figure 2.**
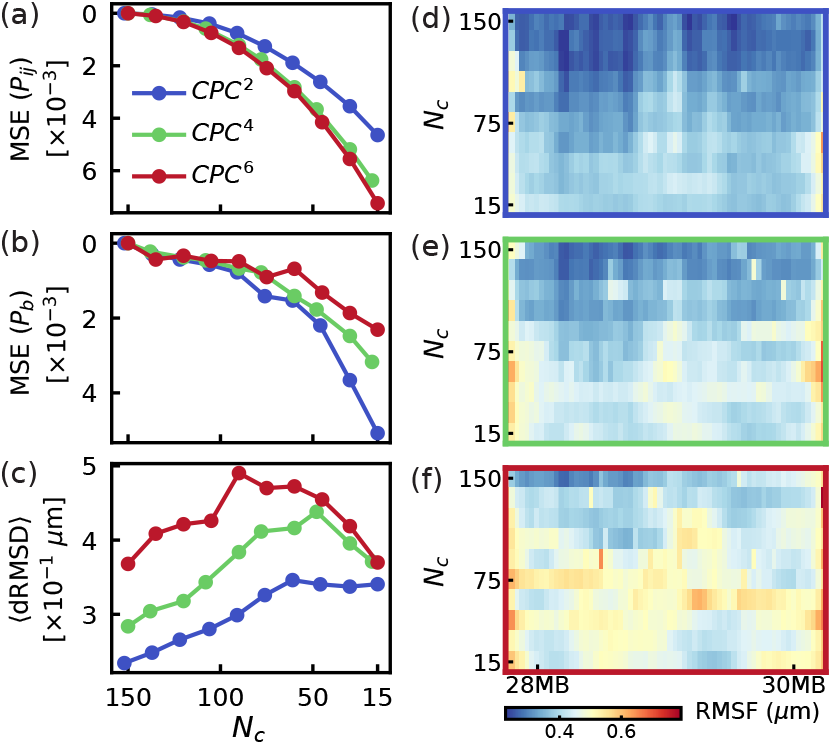
Valency-dependent chromatin stability and mobility under CPC depletion. (a–c) Structural response as the total number of CPC-mediated links *N*_*c*_ is reduced from the saturated reference state (*N*_*c*_ ≃ 150). Shown are the mean-squared error (MSE) of the contact probability map *P*_*ij*_ relative to the reference ensemble (a), the MSE of the boundary-probability profile *P*_*b*_(*i*) relative to the reference ensemble (b), and the ensemble-averaged dRMSD reporting conformational heterogeneity (c). (d–f) Spatial pattern of locus mobility during depletion. Heat maps show the per-locus root-mean-square fluctuation (RMSF) as a function of decreasing *N*_*c*_ for the same valencies shown in (a–c).

In the high-valency limit (*k* = 6), CPCs and chromatin beads co-assemble into pronounced, spatially distinct hubs (Fig. S6). While this topology is less constraint-efficient at global compaction compared to the dispersed *k* = 2 network, it creates a larger pool of residual degrees of freedom and thus a more deformable ensemble. Consistent with this picture, the chromatin conformational heterogeneity increases as *N*_*c*_ is reduced and reaches a maximum at intermediate depletion (Fig.2c), indicating a crossover from an over-constrained state to a regime dominated by generic polymer fluctuations. This regime supports diverse metastable configurations, manifested as spatially localized mobility hotspots (Fig.2f). These trends align with the expectation that multivalent binders can stabilize local clusters while keeping the polymer globally compliant, thereby enhancing structural plasticity [23, 37].

Motivated by the two limiting regimes above, we considered the complex biomolecular component conditions *in vivo* and hypothesized that realistic chromatin folding may require a heterogeneous mixture of CPC valencies. We therefore built mixed-valency hybrid ensembles by continuously titrating the CPC pool from pure *CPC*^2^ to pure *CPC*^6^ while holding the total connectivity budget fixed at *N*_*c*_ ≃ 150 (Fig. 3 and Figs. S7,S8). CPC6-enriched mixtures achieve lower training error (Fig. S7), consistent with hub-forming co-bridging being important for population chromatin contacts. At the same time, the ensemble becomes progressively more heterogeneous, reflecting the lower constraint efficiency of high-*k* hubs. Thus, better Hi-C fidelity obtained by enriching high-valency binders comes with an intrinsic increase in conformational diversity, motivating mixed valencies as a route to balance population-level structure and single-cell variability.

**Figure 3.**
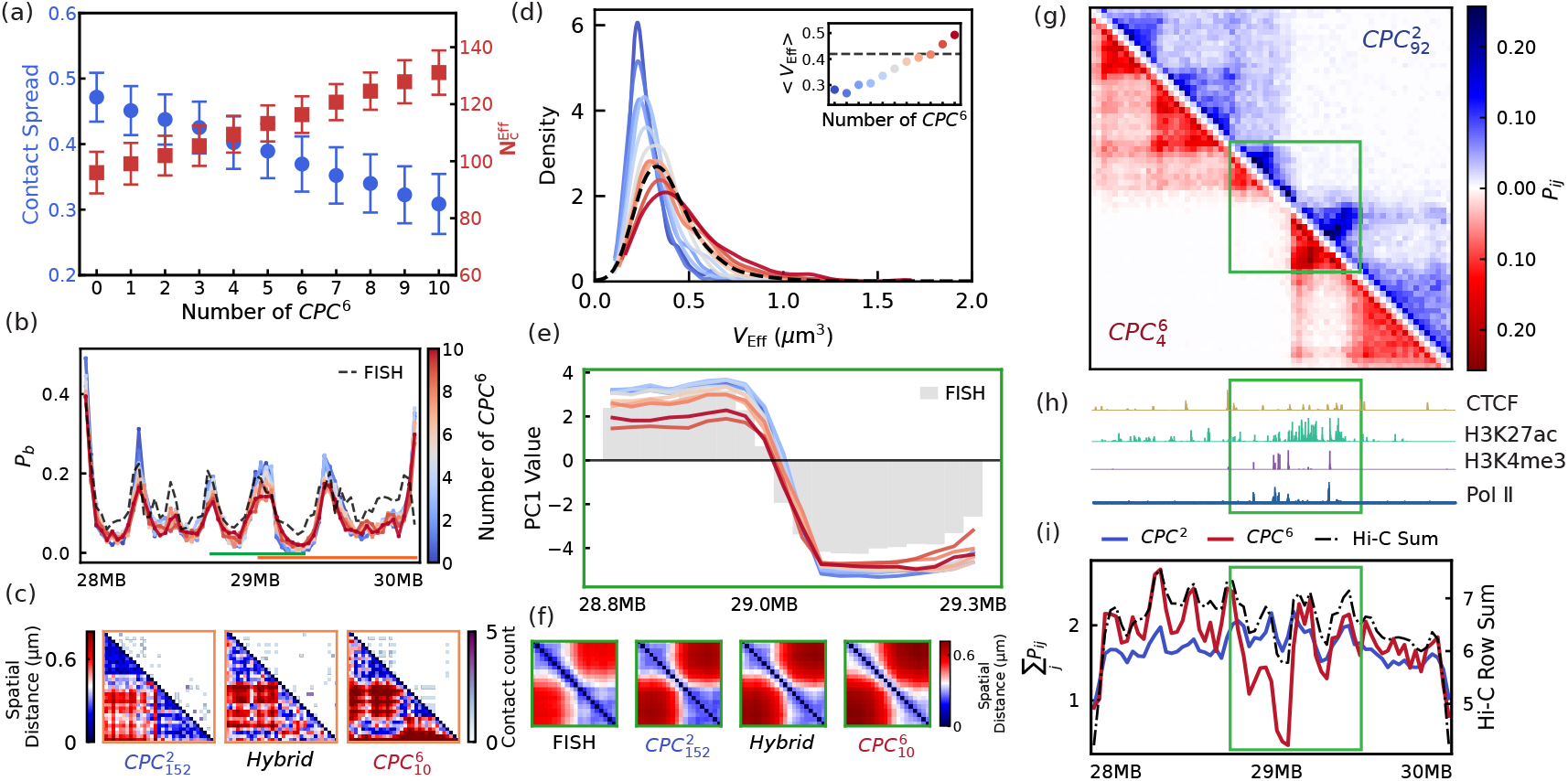
Synergy between low- (*k* = 2) and high-valency (*k* = 6) CPCs in different mixture systems. (a) Contact spread (blue) and total effective number of CPC-mediated connections 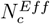 (red) versus mixture composition. Contact spread quantifies how uniformly CPC-mediated connections are distributed across loci (large: broadly distributed; small: hub concentrated). (b) Boundary probability profiles for mixed system with increasing *CPC*^6^ fraction. Orange (29.0–30.0 Mb) and green (28.8–29.3 Mb) markers indicate subregions analyzed in (c) and (e–i), respectively. Dashed curves indicate the experimental data. (c) Representative conformations for pure *CPC*^2^, a mixed system, and pure *CPC*^6^. Top: CPC-mediated connection networks; bottom: corresponding spatial distance matrices. (d) Distributions of effective chromatin volume *V*_*Eff*_ across ensembles. Inset: mean *V*_Eff_ versus mixture composition. Dashed curves indicate the experimental data. (e) PC1 profile of distance matrix for the orange-marked subregion, compared with the experimental range (grey shading). (f) Zoomed spatial distance maps for the green-marked subregion. Decomposition of the mixed-system contact map into CPC-type-specific contributions: contacts mediated by *CPC*^2^ and by *CPC*^6^. (h) Experimental ChIP-seq tracks [36]. (i) Genomic binding profiles of *CPC*^2^ and *CPC*^6^, compared with the experimental Hi-C row sum.

Although increasing the *CPC*^6^ fraction raises the total number of CPC-mediated connections, it simultaneously reduces the contact distribution (quantified by contact spread), implying that connections become more concentrated onto a smaller set of loci (Fig. 3a). Meanwhile, TAD-like insulation decreases with increasing the *CPC*^6^ fraction (Fig. 3b). Representative snapshots illustrate the microscopic origin of this trend (Fig. 3c): the high-valency system concentrates links into discrete, block-like multiway clusters, whereas the low-valency system produces a diffuse network of pairwise bridges that compact the chromatin and contribute to enhancing the domain insulation. In parallel, the effective volume distribution shifts toward larger values as *CPC*^6^ becomes more prevalent (Fig. 3d), consistent with the reduced constraint efficiency of high-valency hubs at fixed *N*_*c*_ (Fig. S3).

At finer resolution, the limitations of the high-valency limit become apparent. In the subregion focusing on the TAD boundary (Fig. 3e), the PC1 profile for pure *CPC*^6^ captures the dominant sign change across the boundary, indicating that hub formation recovers the chromatin compartmentalization. However, it smooths out weaker intra-domain modulations, yielding a PC1 trace that lacks the smaller-amplitude structure present in the experimental range. The zoomed distance maps pinpoint the structural origin (Fig. 3f): pure *CPC*^6^ builds a robust domain scaffold with strong boundary segregation, but leaves sparse patches of weakened connectivity within domains. Reintroducing low-valency *CPC*^2^ bridges preferentially restores short-to intermediate-range contacts, filling these gaps without erasing the hub-defined scaffold. As a result, mixed-valency systems preserve domain-level organization while better reproducing fine-scale boundary.

Decomposing the mixed-system contacts by CPC type makes the division of labor explicit (Fig. 3g). High-valency *CPC*^6^ contributes a sparse set of strong, localized co-bridging events that disproportionately shape domain organization, consistent with hub-like multi-way contact foci. Low-valency *CPC*^2^ instead provides a broadly distributed background of pairwise bridges that densifies the interaction network, promotes uniform compaction, but strengthens boundary insulation. This “hub-and-matrix” architecture is echoed by the epigenomic context (Fig. 3h,i): *CPC*^6^ enrichment peaks near CTCF-associated anchor loci and overlaps regions of strong enhancer activity (H3K27ac), consistent with higher-order anchor and enhancer assemblies beyond isolated pairwise loops [38–41], whereas, in addition to widespread general distribution, *CPC*^2^ enrichment shows promoter-proximal features (H3K4me3 and Pol II), consistent with abundant low-order bridging that stabilizes local regulatory neighborhoods [42, 43].

To probe causality at anchors, we performed *in silico* CTCF perturbation simulations by setting the learned affinities {*α*_*ij*_} involving loci with strong CTCF signal to zero, thereby removing CTCF-linked interactions from the affinity landscape. This perturbation reduces boundary insulation, with the effect most pronounced in the pure*CPC*^2^ system, while a *CPC*^6^-dominated system shows a more moderate change in global connectivity (Fig. S9). The differential sensitivity supports a valency-based “cocktail” mechanism in which high-valency CPCs stabilize sparse anchor/enhancer-centered hubs that drive domain-scale features and heterogeneity, whereas low-valency CPCs form a matrix of constraint-efficient pairwise links refining insulation. These roles have plausible biological counterparts: hub-like behavior aligns with multicomponent condensates at super-enhancers (Mediator/BRD4-rich) and multivalent heterochromatin effectors such as HP1*α* [10, 44], as well as higher-order clustering of architectural anchors [40, 41]; the matrix-like component is consistent with predominantly pairwise bridging mechanisms such as YY1-mediated enhancer-promoter looping, LDB1-linked assemblies and CTCF-cohesin loops [45–47].

To further disentangle locus specificity from constraint topology, we carried out additional simulations in which the learned, locus-resolved affinity landscape {*α*_*ij*_} was replaced by a non-specific “uniform-*α*” baseline that depends only on genomic separation (Fig. S10). This model preserves generic polymer scaling but removes the positional information needed to encode locus-specific architectural elements. Consequently, the resulting ensembles fail to recover TAD/loop-like features and exhibit conformational heterogeneity far above the experimental reference for all *k*. These results propose a clear division of roles: the structured, data-inferred {*α*_*ij*_} provides the locus-specific interaction blueprint that confines the chromatin ensemble and specifies where architectural organization is favored [29], whereas CPC valency governs how that blueprint is realized by tuning the topology of constraints and the structural plasticity within the confined chromatin conformational manifold.

Finally, our results suggest a two-handle control strategy for 3D genome folding that separates where interactions are favored from how they are executed in space. First, sequence- and epigenome-defined chromatin states establish locus-resolved interaction propensities. In CPC-MEM, this blueprint is encoded by the inferred affinity landscape {*α*_*ij*_}, consistent with the view that epigenetic state and physicochemical interactions jointly constrain chromatin folding and can reinforce cell-type-specific organization and memory [43, 48, 49]. Second, the nuclear milieu provides an orthogonal systems-level knob: a diverse repertoire of nuclear condensates and bodies, differing in composition, stoichiometry, and material properties, naturally realizes different effective multivalent interaction architectures and exchange kinetics, thereby tuning how efficiently and how dynamically the same locus-specific blueprint is implemented [50–52]. In this view, CPC-MEM offers a unifying, quantitative route to integrate these layers within a single data-constrained ensemble framework: {*α*_*ij*_} specifies locus selectivity, while condensate or CPC valency (and abundance) regulates the compaction-plasticity balance and the efficiency of executing that blueprint in 3D genome folding.

Looking forward, CPC-MEM offers a natural launching point for active genome folding. ATP-driven loop extrusion and transcription-coupled remodeling can continuously reshuffle chromatin contacts in time while largely preserving locus-specific interaction preferences [22, 53–55]. Embedding the nonequilibrium effects into CPC-MEM can turn the framework into a predictive platform for when activity reorganizes domain and modulates cell-to-cell variability under a fixed genomic blueprint [56]. We anticipate that such extensions will enable quantitative, parameter-matched comparisons to acute perturbations and time-resolved single-cell measurements of genome architecture and dynamics [22].

## Acknowledgments

X.C. thanks the support from the National Natural Science Foundation of China (Grant Nos. 12474201 and 32201020), the General Program of the Guangdong Basic and Applied Basic Research Foundation (Grant No. 2024A1515010862), the Guangdong Provincial Project (Grant No. 2023QN10X037) and the Guangdong S&T Program (Grant No. 2025A0505000027). The authors also acknowledge the Green e Materials Laboratory (GeM) and HPC+AI Intelligence Computing Center at the Hong Kong University of Science and Technology (Guangzhou) for providing computational support.

## Supplemental Material

## S1 Studied genomic region and experimental datasets

We focus on a 2 Mb segment on human chromosome 21 in IMR90 cells. Unless otherwise stated, we use hg38 coordinates chr21:28,000,071–29,949,939 (corresponding to hg19 chr21:29,372,390–31,322,257). The chromatin polymer is coarse-grained at 30 kb resolution, yielding *N* = 65 beads, each representing a genomic locus. Population-averaged Hi-C contact information for IMR90 cells is taken from Refs. [3, 16] for the locus above. We extract the corresponding contact map, then aggregate contacts to 30 kb bins, and finally normalize it to obtain target contact probabilities *P*_*ij*_ used for maximum-entropy inference [29]. Independent single-cell structural validation is based on super-resolution single-cell chromatin tracing measurements for the same IMR90 locus [16]. These data provide 3D coordinates for the traced loci in many individual cells, from which we calculate pairwise spatial distance distributions for selected locus pairs and single-cell distance matrices and structural statistics presented in Fig. 1 and Fig. S1.

Hi-C reports an population-averaged contact probability, whereas imaging reports single-cell spatial distances. As chromatin conformations are heterogeneous, there is generally no one-to-one mapping between contact probability and mean distance. This mismatch underlies the “Hi-C–FISH paradox” summarized in Fig. S1 [17, 18]. To generate polymer ensembles to both Hi-C and imaging on a consistent footing, we characterize the empirical relationship between Hi-C contact probabilities *P*_*ij*_ and imaging-derived mean spatial distances *r*_*ij*_ (Fig. S1c). However, a simple power-law mapping, *P* ∝ *r*^−*β*^, fits the short-distance regime but deviates at larger separations (Fig. S1d).

We therefore use a sigmoidal (tanh) function that remains well-behaved across the full dynamic range and directly defines a smooth contact observable from distances:

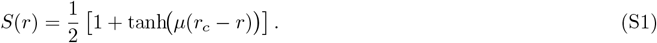

Here *r*_*c*_ is an effective contact range and *µ* controls the sharpness of the switch. We fit (*r*_*c*_, *µ*) using the empirical Hi-C-FISH relationship (Fig. S1e). As shown in Fig. S1f, this tanh-based calibration better preserves the observed contact-probability decay with genomic separation compared to a global power-law. Thus, it is used for the following contact probability modeling.

## S2 Models and simulation methods

### S2.1 Maximum-entropy principle model

We reconstruct chromatin structural ensembles using a maximum-entropy principle polymer model in which the least-biased distribution consistent with experimental Hi-C contact statistics is realized by an effective energy functional [29]. In both the conventional CC-MEM and our CPC-MEM, the total potential energy takes the same form

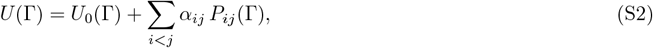

where Γ denotes the system configuration, *U*_0_ is a reference potential encoding baseline polymer physics, {*α*_*ij*_} are Lagrange multipliers inferred from Hi-C, and *P*_*ij*_(Γ) is the model-specific contact observable. The two frameworks differ in the microscopic definition of *P*_*ij*_: CC-MEM treats contacts as direct chromatin-chromatin proximity, whereas CPC-MEM treats contacts as CPC-mediated bridging events under an explicit valency constraint.

### S2.2 Coarse-grained chromatin polymer model

We model the chromatin as a coarse-grained semiflexible polymer of *N* beads. All simulations use reduced units with energy unit of 1.0 and length unit of 1.0. By matching the distance between adjacent beads in our polymer model to experimental imaging measurements, the length unit is estimated to correspond to approximately 133 nm in physical space. Unless otherwise stated, parameter values below are identical for CC-MEM and CPC-MEM.

Consecutive beads are connected by a Morse bond potential,

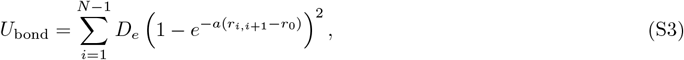

where *r*_0_ = 1.4 is the equilibrium bond length, *D*_*e*_ = 20.0 is the well depth, and *a* is the inverse-length range parameter that sets the bond curvature (stiffness) near *r*_0_. For small fluctuations about *r*_0_, the Morse potential is locally harmonic with effective spring constant *k*_bond_ = 2*D*_*e*_*a*^2^. We therefore choose 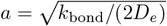 with the local stiffness *k*_bond_ = 10.0.

Chain semiflexibility is enforced by an angular cosine potential,

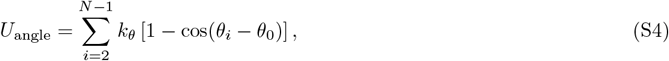

where *θ*_*i*_ is the angle between adjacent bonds, *θ*_0_ = *π/*2, and *k*_*θ*_ = 0.2.

Nonbonded steric repulsion between chromatin beads (for |*i* − *j*| ≥ 2) is modeled by a truncated-and-shifted Lennard-Jones (WCA) potential,

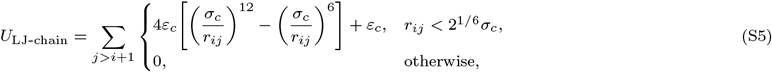

with *ε*_*c*_ = 2.1 and *σ*_*c*_ = 0.5.

To mimic nuclear confinement, all beads are contained in a spherical cavity of radius *R*_w_ = 10.0, implemented via a repulsive WCA wall potential acting on the distance to the boundary 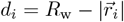,

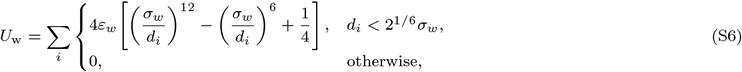

with *ε*_*w*_ = 1.0 and *σ*_*w*_ = 1.0.

### S2.3 Specific interaction term: CC-MEM versus CPC-MEM

The second term in Eq. (S2) encodes the Hi-C constraints through {*α*_*ij*_} and the contact observable *P*_*ij*_(Γ). We use a smooth contact switch

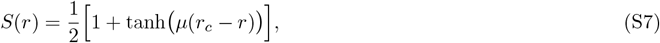

which transitions from ≈ 1 (contact) to ≈ 0 (no contact) around *r*_*c*_, with steepness *µ*. Unless otherwise noted, *µ* = 14.0 and *r*_*c*_ = 1.2.

In CC-MEM, contacts arise from direct spatial proximity between chromatin beads, so

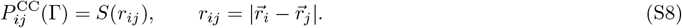

With this definition, *α*_*ij*_ acts as an effective direct attraction (or repulsion) between loci *i* and *j*.

In CPC-MEM, we introduce a finite pool of *M* explicit CPCs, each represented as a diffusing spherical bead with excluded-volume interactions (WCA form) against both chromatin beads and other CPCs (parameters *ε*_*p*_ = 2.1, *σ*_*p*_ = 1.5). Now, the configuration is 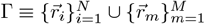.

Each CPC *m* is assigned an interaction valency *k*, meaning it can simultaneously engage at most *k* chromatin loci. Operationally, for a given configuration Γ, we define 𝒩_*m*_ as the set of the *k* nearest chromatin beads to CPC *m* (optionally restricted to beads within a maximum capture distance *r*_max_ = 1.5). The contribution of CPC *m* to bridging loci *i* and *j* is

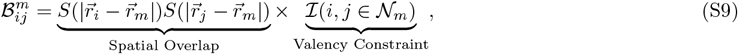

where indicator ℐ enforces the valency constraint.

We define the locus-locus contact observable as the probability that *i* and *j* are bridged by at least one CPC in the current configuration. By assuming independent bridging attempts across CPCs, this gives

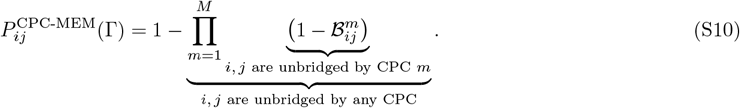

With this definition, *α*_*ij*_ no longer prescribes a direct bead-bead attraction. Instead, it quantifies the propensity for loci *i* and *j* to be bridged by a shared CPC under an explicit, valency-controlled binding topology.

### S2.4 Monte Carlo sampling and maximum-entropy optimization

For a fixed parameter set {*α*_*ij*_}, we sample the joint chromatin-CPC configuration from the Boltzmann distribution associated with the total potential *U*(Γ). Sampling is performed with a Metropolis-Hastings Monte Carlo (MC) scheme in a cubic box of side length *L* = 20 under spherical confinement [35]. Unless stated otherwise, one optimization round consists of 2 × 10^7^ attempted MC moves. The first 50% are discarded as equilibration step and the remaining configurations are recorded every 2 × 10^4^ attempts to estimate ensemble averages.

To efficiently generate polymer conformations while maintaining connectivity, we combine local and global chain updates with CPC diffusion moves. At each attempted step we select one move type with a prescribed relative frequency (weights below), propose an update, and accept/reject using the standard Metropolis criterion. The move set includes: (i) local single-bead displacements for chromatin beads (weight 100); (ii) pivot rotations (weight 20); (iii) crankshaft/double-pivot updates for local reorientation (weight 20); (iv) CPC translation/reptation-like diffusion moves to enhance binder mobility (weight 5); (v) rare loop/topology-accelerating moves used to reduce long-lived entanglement (weight 1). We verified that observables reported are stable with respect to reasonable variations of these move frequencies.

Given the sampled ensemble at round *t*, we compute the model-predicted contact observables 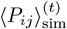 by averaging *P*_*ij*_(Γ) over recorded snapshots. For CPC-MEM, *P*_*ij*_ is evaluated from the binder-mediated bridging definition (Eq. S10), whereas for CC-MEM it reduces to the proximity-based definition *S*(*r*_*ij*_) (Eq. S7). All reported contact maps and derived quantities are calculated from these ensembles.

The interaction parameters *α*_*ij*_ are inferred by iteratively matching ⟨*P*_*ij*_⟩_sim_ to the experimental Hi-C targets 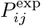. We update *α*_*ij*_ using a stochastic gradient step with a decaying learning rate,

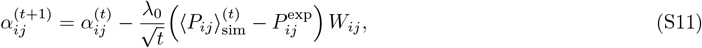

with *λ*_0_ = 2.0. To emphasize long-range, low-probability contacts during early learning, we use a distance-dependent weight *W*_*ij*_ = | *i ™ j* | ^*γ*^ with *γ* = 0.8. The matrix {*α*_*ij*_} is kept symmetric by construction, and we monitor convergence using the MSE between simulated and experimental contact probabilities (Fig. S2). To reduce estimator noise in ⟨*P*_*ij*_⟩_sim_ during learning, each round is executed as multiple independent MC replicas (10 cores in parallel in our implementation). Replica averages are combined to obtain 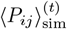 for the update in Eq. (S11).

Evaluating the many-body CPC-mediated contact observable, 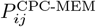, can suffer from underflow when the number CPCs is large and when many factors are close to unity. We therefore calculate this term in a numerically stable form using log-domain accumulation (a log-sum-exp evaluation), i.e., we accumulate 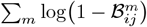 and then exponentiate once to obtain the product.

To enforce the valency constraint efficiently, we note that each CPC only needs information about its *k* nearest chromatin beads (within the interaction range). For each CPC, we maintain a fixed-size list of the *k* closest beads and update it by a partial-selection scan. This avoids repeated global sorting and reduces the per-update cost from generic neighbor-search scaling to an *O*(*Nk*) scan per CPC (with small *k* ≤ 6 in this work). We implement valency-specific, allocation-free kernels for *k* = 2–6, so that the bridging scores and contact observables are evaluated without heap allocations in the inner loop.

Simulation and optimization codes used in this work are available at https://github.com/CodePioneer42/PolymermcCPCMEM.

### S2.5 Validation of maximum-entropy optimization

To assess validation of the iterative maximum-entropy inference, we monitor the evolution of the ensemble generated by a representative system 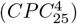 throughout the optimization (Fig. S2). The procedure starts from {*α*_*ij*_} = 0, for which the polymer behaves as a confined semiflexible chain and the contact map exhibits only generic distance-dependent decay. After the first few parameter updates, the simulated contact map rapidly develops the dominant experimental architectural features (Fig. S2a), accompanied by a sharp decrease of the loss (MSE; Fig. S2b).

We interpret the subsequent iterations as a refinement regime: global agreement saturates as Pearson correlation coefficient (*pcc*) reaches *r* ≃ 0.99 by late rounds, while the inferred interaction landscape becomes progressively more structured. This sharpening is reflected by the growth of the parameter variance (Fig. S2c), indicating increased contrast between high-affinity and background interactions required to match locus-resolved contact heterogeneity. In parallel, ensemble-level polymer observables exhibit physically consistent stabilization: locus-wise fluctuations (RMSF) are reduced from the initial homopolymer baseline and then plateau as the inferred constraints converge (Fig. S2d) and the radius of gyration decreases toward a compact steady state (Fig. S2e). Together, these characteristics indicate that the optimization reaches a stable fixed point in both the inferred {*α*_*ij*_} and the resulting structural ensemble.

## S3 Definitions of physical quantities

To characterize the structural ensembles produced by CC-MEM and CPC-MEM and to compare them with Hi-C and imaging measurements, we define the following observables. Throughout, Γ denotes a single configuration (snapshot) of the simulated system, and angle brackets ⟨·⟩ denote an average over the converged ensemble (MC samples after equilibration).

### S3.1 Contacts and connectivity

In both CC-MEM and CPC-MEM, the model defines an instantaneous contact observable *P*_*ij*_(Γ) ∈ [0, 1]. The corresponding contact probability map is the ensemble average

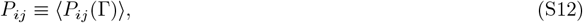

which is the quantity fitted to (or compared against) the experimental Hi-C contact probabilities.

We calculate the standard contact-decay curve as a function of genomic separation *s* = |*i* − *j*|,

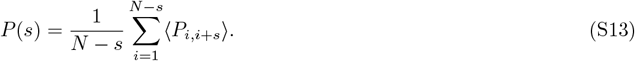

When useful, an effective decay exponent is obtained from a log–log fit over a specified *s* range.

To describe the distribution of realized CPC-mediated links, we construct a per-snapshot connection (link) map *L*_*ij*_(Γ) ∈ {0, 1}, where *L*_*ij*_(Γ) = 1 if there exists at least one CPC *m* that simultaneously engages loci *i* and *j* under the same rule used for the connection snapshots (i.e., both loci are within the CPC interaction range and satisfy the valency selection via membership in 𝒩_*m*_); otherwise *L*_*ij*_(Γ) = 0.

For a system with *M* CPCs of interaction valency *k*, we define a connectivity budget *N*_*c*_ as the maximum number of pairwise links that would be available if each CPC simultaneously engaged *k* loci and thereby formed a *k*-clique:

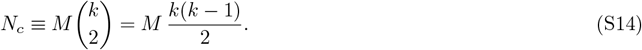

This budget depends only on (*M, k*) and is the quantity we match across valencies when we set *N*_*c*_ ≃ 150 by adjusting *M* accordingly. In depletion/titration protocols, the control parameter is *N*_*c*_.

For each configuration Γ, we define the realized (effective) CPC-mediated connectivity as the number of distinct locus pairs connected by at least one CPC under the same rule used to generate the connection snapshots, and the ensemble-averaged effective connectivity is calculated as

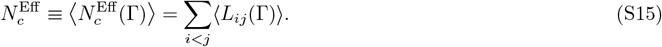

We define the integer connection count as

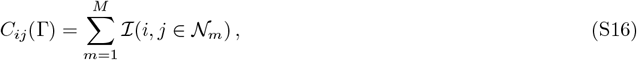

which records how many CPCs simultaneously connect the same pair (*i, j*) in a configuration. We use *L*_*ij*_ when we need a binary link map and *C*_*ij*_ when we need multiplicity information (e.g., contact spread and CPC-type decomposition).

The contact spread is calculated from the ensemble-averaged connection-count matrix ⟨*C*_*ij*_⟩. We coarse-grain ⟨*C*_*ij*_⟩ into non-overlapping 4 × 4 blocks 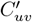 in (*u, v*) space and define a block-occupancy indicator

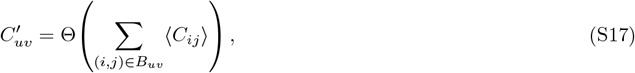

where Θ(·) is the Heaviside step function. The contact spread is then calculated as

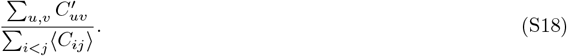

Smaller contact spread indicates that CPC-mediated links are concentrated into fewer genomic neighborhoods (hub-dominated patterns), whereas larger contact spread indicates more spatially distributed connectivity.

Local connectivity along the polymer is summarized by the per-locus link number

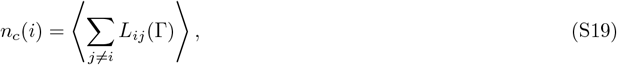

which measures how the CPC-mediated link network is distributed across genomic positions.

### S3.2 Structural ensemble analysis

We quantify global compaction using the radius of gyration,

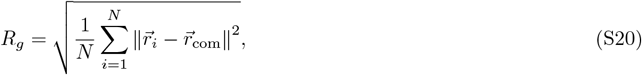

Where 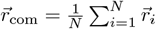 is the center-of-mass coordinate of the chromatin beads. As a complementary, shape-aware measure that does not assume spherical symmetry, we define an effective volume *V*_eff_ as the volume of the Convex Hull of the bead coordinates 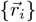.

To compare conformations without performing any rigid-body alignment, we use the distance root-mean-square deviation (dRMSD), which measures the discrepancy between two inter-bead distance matrices. For two conformations *ν* and *ξ*, let 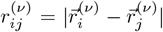 denote the distance between beads *i* and *j* in conformation *ν* (and similarly for *ξ*). When experimental traces have missing loci, we restrict the comparison to a “core” set of beads detected in more than 95% of experimental structures; the core size is denoted *N*_core_. The dRMSD is then

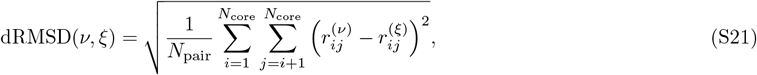

where 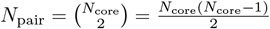 is the number of calculating pairs in the core set. We use intra-ensemble dRMSD statistics (pairwise dRMSD among conformations drawn from the same ensemble) to quantify structural heterogeneity, and inter-ensemble dRMSD statistics (between simulated and experimental conformations) to quantify agreement with experiments.

To visualize the resulting high-dimensional conformational relationships, we perform metric multidimensional scaling (MDS) on the dRMSD distance matrix. Given dissimilarity *δ*_*νξ*_ = dRMSD(*ν, ξ*), MDS places conformations as points in 2D such that Euclidean distances *d*_*νξ*_ approximate *δ*_*νξ*_. We quantify embedding quality by the normalized stress function,

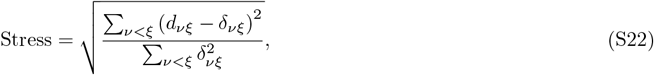

following standard MDS practice.

For selected subregions, we further characterize dominant modes of structural variability by principal-component analysis. Specifically, we vectorize the upper-triangular entries of the corresponding subregion distance matrices (or, equivalently, the distance vectors) across the ensemble and compute the first principal component (PC1) to obtain a one-dimensional summary of the leading structural variation within that subregion.

### S3.3 Domain boundaries and model-validation metrics

We quantify domain insulation using a distance-based separation-score analysis computed on individual 3D conformations. For each conformation, the separation-score profile is used to call stochastic domain boundaries, and the boundary occurrence probability is obtained by averaging the per-structure calls over the ensemble. The same pipeline is applied to experimental single-cell traces and to simulated structures.

For a given conformation Γ with bead coordinates 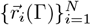, we define the pairwise distance matrix *D*(Γ) with element (*u, v*)

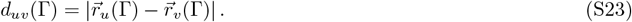

For each candidate boundary between beads *i* and *i* + 1, we consider an upstream window 𝒰 (*i*) = {*i* − ω, …, *i* − 1} and a downstream window 𝒱 (*i*) = {*i* + 1, …, *i* + *ω*}, where *ω* is the window size in beads (*ω* = 6, corresponding to 180 kb at 30 kb resolution). Let *N*_𝒰_ (i; Γ) and *N*_𝒱_ (*i*; Γ) denote the numbers of valid (non-missing) beads in the two windows for conformation Γ. The per-structure separation score is defined as the mean cross-window distance,

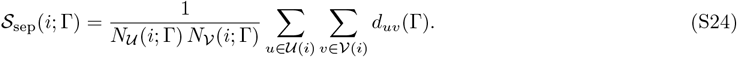

A larger 𝒮_sep_(*i*; Γ) indicates that the two flanking segments are, on average, farther apart in 3D space, consistent with stronger local insulation at locus *i*.

To identify discrete boundaries in a single conformation, we perform peak calling on the profile 𝒮_sep_(*i*; Γ) as a function of *i*. A locus *i* is labeled as a boundary in conformation Γ if it is a local maximum with prominence exceeding 5% of the dynamic range of 𝒮_sep_(·; Γ) in that conformation. We encode this as a binary indicator variable *b*(*i*; Γ) ∈ {0, 1}.

The ensemble-averaged boundary probability is then defined as the frequency of boundary occurrence at each locus,

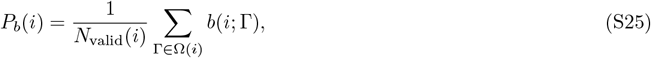

where Ω(*i*) is the set of conformations for which both windows contain at least one valid bead (so that 𝒮_sep_(*i*; Γ) is defined), and *N*_valid_(*i*) = |Ω(*i*)|. For experimental traces, beads with missing coordinates are excluded from all distance evaluations; if either window contains no valid beads for a given (*i*, Γ), that entry is omitted from Ω(*i*).

To quantify deviations from a reference under titration/depletion, we report mean-squared errors (MSE). For two symmetric matrices *A*_*ij*_ and *B*_*ij*_ (e.g., ensemble distance or contact maps):

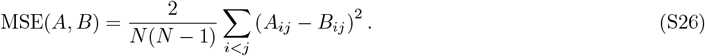

For one-dimensional profiles *x*(*i*) and *y*(*i*) defined on the same genomic bins (e.g., the boundary probability *P*_*b*_(*i*)):

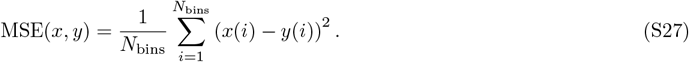

In Fig. 2a we use MSE_2D_ between the depleted contact map and the saturated reference map; in Fig. 2b we apply MSE_1D_ to the corresponding boundary probability profile *P*_*b*_(*i*).

## S4 Theoretical Derivation: Valency-Dependent Structural Heterogeneity

Here we provide a constraint-counting rationale for why, under the constant-*N*_*c*_ protocol used in the main text, increasing CPC valency *k* tends to increase ensemble heterogeneity. The core point is that a fixed pairwise-link budget *N*_*c*_ does not fix the number of independent geometric constraints imposed on the polymer: multivalent CPCs can spend many links to build locally redundant hubs, thereby leaving more residual degrees of freedom for the chromatin backbone [57–59].

### S4.1 Constant connectivity budget

In the constant-*N*_*c*_ comparisons, we fix the total number of CPC-mediated pairwise links (the connectivity budget) to

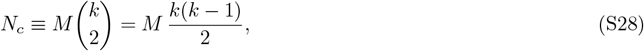

where *M* is the number of CPCs and *k* is the maximum number of loci that a CPC can engage simultaneously (i.e., a fully occupied CPC induces a *k*-clique of pairwise links). Thus, at fixed *N*_*c*_,

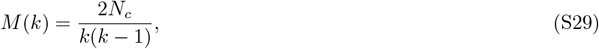

so increasing valency necessarily reduces the number of CPCs available to distribute the same overall link budget.

### S4.2 Independent geometric constraints per CPC

A CPC that simultaneously engages *k* loci can be viewed, in a Maxwell-counting sense, as forming a local rigid cluster whose internal geometry is fixed up to global rigid-body motions [57–59]. In 3D, the maximum number of independent relative constraints associated with such a rigid cluster scales as 3*k* − 6 for *k* ≥ 3 [57–59]. However, achieving this local rigidification consumes 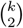 pairwise links from the global budget. This motivates the constraint efficiency

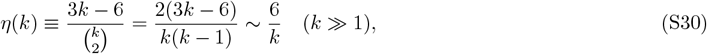

which decreases with increasing *k*. Intuitively, high-valency CPCs allocate many links to impose constraints within the hub, but yield fewer independent constraints per link than predominantly pairwise bridging.

In practice, not all nominal constraints are independent: cycles in the interaction graph generate redundancy and reduce the effective constraint count [58, 59]. We therefore write the effective number of independent constraints contributed by CPC-mediated topology as

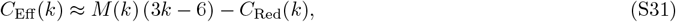

where *C*_Red_(*k*) ≥ 0 summarizes redundant constraints. For the scaling argument below, the key feature is that the leading term *M*(*k*)(3*k* − 6) ∝ *N*_*c*_ *η*(*k*) decreases with *k*, and redundancy can only further decrease *C*_Eff_.

### S4.3 Degrees of freedom and heterogeneity

Let *F*(*k*) denote the number of accessible degrees of freedom after constraints are imposed. Baseline polymer constraints from chain connectivity, bending rigidity, excluded volume, and confinement are identical across valencies and can be absorbed into a constant *F*_0_. The valency dependence then enters through *C*_Eff_ (*k*):

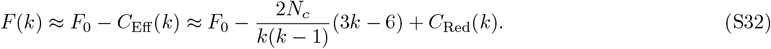

Using Eq. (S30), the leading trend can be summarized as

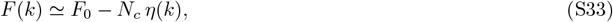

so that, at fixed *N*_*c*_, larger *k* leaves more accessible degrees of freedom because the independent constraints per link are less efficient.

Finally, under a small-fluctuation (harmonic) picture, the typical amplitude of structural fluctuations increases with the square root of the number of accessible modes [60]:

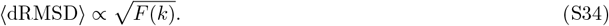

Eqs. (S33)–(S34) therefore predict that, under a fixed connectivity budget *N*_*c*_, increasing valency increases ensemble heterogeneity, consistent with the trends observed in Fig. S3.

## S5 Fitting results across CPC valency

This section compares CPC-MEM ensembles across interaction valencies *k* = 2 to 6 under two complementary protocols. In the first protocol we match the connectivity budget *N*_*c*_ in Eq. (S14) across valencies so that the maximum available number of CPC-mediated pairwise links is approximately fixed (*N*_*c*_ ≃ 150). In the second protocol we fix the CPC number *M* and allow *N*_*c*_ to vary with *k*. Together, these two settings separate effects arising from the total link capacity from effects arising from the local topology imposed by CPC valency.

### S5.1 Scenario 1: Constant connectivity budget (*N*_*c*_ ≃ 150Nc150)

Fig. S3 summarizes the constant-*N*_*c*_ comparison. Because the maximum link capacity is matched across *k*, all systems reproduce the population-averaged Hi-C map with similarly high rank agreement (Fig. S3a), and the corresponding *P*(*s*) decay profiles are also comparable (Fig. S3c). This indicates that, at the level of ensemble-averaged contact statistics, fixing *N*_*c*_ largely controls how much long-range contact probability the model can allocate.

Despite the similar map-level agreement, the implied single-cell geometry is not equivalent across valencies. The mean spatial-distance matrices (Fig. S3b) show that low valency (e.g., 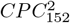) produces sharper, more localized loop-like features, consistent with a more rigid and over-regularized realization of the same Hi-C constraints. As *k* increases, these localized distance features broaden and become more diffuse, reflecting increased structural plasticity and a larger family of geometries compatible with the population contact map.

Single-cell validation against imaging further highlights an intermediate-valency optimum. For the locus-pair distance distributions (Fig. S3d), *k* = 4 best captures both peak positions and distribution widths across the tested probe pairs, whereas low *k* tends to over-constrain distances and high *k* tends to over-broaden them. Consistent with this, the boundary probability profiles (Fig. S3e) reveal a characteristic low-valency artifact: 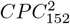 (and, to a lesser extent, the lowest-*k* intermediate case) exhibits unusually sharp, high-amplitude peaks compared with experiment, indicating boundaries that are overly persistent across snapshots. Intermediate valency yields a smoother boundary-probability landscape that better resembles the experimentally inferred variability.

The compaction and network-usage metrics further separate the valency groups. Under fixed *N*_*c*_, increasing *k* necessarily reduces the number of CPCs *M*, which changes how uniformly links can be distributed along the chain. This is reflected in the contacts-per-bead profiles (Fig. S3f), where the highest-valency case 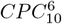 shows a more uneven and less uniformly supported connectivity pattern than lower-*k* systems. At the ensemble level, this redistribution of constraints manifests as increasing structural variability: the dRMSD distributions (Fig. S3g) broaden systematically with *k*, with *k* = 4 again providing the closest match to the experimental heterogeneity benchmark. The same trend is echoed by the effective-volume distributions (Fig. S3h), which become broader and shift toward larger values at high valency, consistent with more expanded and heterogeneous conformations.

Overall, at fixed connectivity budget *N*_*c*_, the main distinction across valency is therefore not whether Hi-C can be fitted, but how the same Hi-C constraints are realized as single-cell geometry, boundary persistence, local compaction, and ensemble heterogeneity.

### S5.2 Scenario 2: Constant number of CPCs (*M* = 40M40)

Fig. S4 compares systems at fixed CPC abundance (*M* = 40). In this setting the maximum link capacity 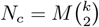 grows rapidly with valency, from *N*_*c*_ = 40 at *k* = 2 to *N*_*c*_ = 600 at *k* = 6; the comparison therefore isolates how interaction valency controls architectural capacity when CPC concentration is held fixed.

The fitted contact maps immediately show that the low-valency system 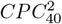 is connectivity-limited: with only *N*_*c*_ = 40 available links, the inferred map is sparse and underestimates long-range contacts, leading to visibly worse agreement with experiment than the higher-*k* cases (Fig. S4a). This limitation is also reflected in the decay curve *P*(*s*), where 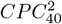 falls below the experimental profile over essentially the full range of genomic separations, whereas increasing *k* progressively restores the correct scaling and long-range contact probability (Fig. S4c).

The corresponding mean spatial-distance matrices show that once *k* ≥ 3 the models recover the overall distance organization much more faithfully, while the low-*k* limit remains biased because the available links are insufficient to stabilize the locus-resolved architecture implied by Hi-C (Fig. S4b). Independent single-cell distance distributions further separate valencies (Fig. S4d): intermediate valency (notably *k* = 4) best captures both peak positions and distribution widths across the tested probe pairs, whereas low *k* tends to over-regularize distances and high *k* tends to over-broaden them. Consistently, boundary statistics indicate that low valency yields overly persistent, sharp boundary peaks, while intermediate *k* reproduces a smoother, more variable boundary landscape (Fig. S4e).

Network-level compaction measures clarify the origin of these trends. The per-locus contact number *n*_*c*_ is strongly suppressed for 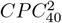 because the total link capacity is too small, while increasing *k* raises *n*_*c*_ toward the experimental reference, with intermediate valency again closest overall (Fig. S4f). In parallel, ensemble heterogeneity increases with valency: the dRMSD distributions broaden systematically with *k*, and intermediate *k* best matches the experimental heterogeneity benchmark (Fig. S4g). The effective-volume distributions show the corresponding compaction shift: the low-valency limit remains too expanded, whereas high valency drives over-compaction relative to experiment; intermediate valency best balances these effects (Fig. S4h).

Together, the fixed-*M* comparison emphasizes that valency is not interchangeable with CPC concentration. At fixed mediator abundance, low valency becomes link-capacity limited and cannot fit Hi-C, whereas high valency can fit Hi-C but tends to overshoot compaction/heterogeneity signatures. The intermediate-*k* regime provides the best simultaneous agreement with both population-averaged and single-cell constraints.

## S6 Perturbation simulations and structural evolution

To test the rigidity-based interpretation in a controlled setting, we carried out an in silico CPC-depletion (titration) protocol in which the learned affinity landscape is held fixed while the number of mediators is reduced. The procedure has two stages: (i) Saturated-state training. For each valency type (*k* = 2, 4, 6), we first optimized the affinity parameters {*α*_*ij*_} at the saturated reference condition, where the connectivity budget was matched across valencies at *N*_*c*_ ≃ 150 (Fig. S3). This produces three reference ensembles that represent the same target chromatin folded state under different CPC valencies. (ii) CPC-depletion at fixed {*α*_*ij*_}. Starting from each saturated reference model, we then decreased the number of CPCs *M* while keeping {*α*_*ij*_} unchanged, and re-equilibrated the system to obtain the CPC-depleted chromatin ensembles. To enable a direct topological comparison across *k* at each CPC-depletion level, we aligned titration steps by matching the corresponding connectivity budget

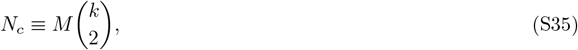

i.e., we chose *M* for each *k* so that all models share the same *N*_*c*_ at a given step. The resulting schedule is listed in Table S1. The structural response to depletion is summarized in Fig. S5.

Across depletion, the three valency classes exhibit distinct failure modes that mirror their different constraint topologies. Global size measures show that the *k* = 2 system remains comparatively compact and changes smoothly as *N*_*c*_ is reduced (Fig. S5c), consistent with constraints being distributed broadly along the polymer. In the same regime, structural deviations from the saturated reference accumulate in a spatially uniform manner (Fig. S5a), indicating a largely homogeneous weakening of the folding scaffold. By contrast, the *k* = 6 system stays more expanded and displays substantially larger fluctuations at matched *N*_*c*_ (Fig. S5c). The corresponding dRMSD distributions broaden markedly and develop heavier tails under depletion (Fig. S5d), consistent with hub-like constraints leaving more residual degrees of freedom even when the nominal link budget is comparable.

Boundary-level results reveal an accompanying robustness-plasticity trade-off. Boundary probability profiles indicate that low-valency systems can support strong, localized insulation features at the saturated state, but these features deteriorate rapidly once links are removed (Fig. S5e). In contrast, high-valency systems display more moderate boundary signatures that decay gradually with depletion, reflecting a more evenly distributed constraint network. Together, these depletion responses provide a direct dynamical complement to the static comparisons in the main text: low valency emphasizes globally distributed, resilient constrainting, whereas high valency concentrates constraints into hubs that promote pronounced, but more fragile and heterogeneous, organization.

**Table S1.** CPC-depletion detailing the number of CPCs (*M*) for each valency group. Steps are aligned by the approximate (*N*_*c*_) to allow for a direct comparison.

| Step | $N_c$ | $CPC^6$ | $CPC^4$ | $CPC^2$ |
| --- | --- | --- | --- | --- |
| 0 (Sat.) | $\sim 150$ | 10 | 25 | 152 |
| 1 | $\sim 135$ | 9 | 23 | 137 |
| 2 | $\sim 120$ | 8 | 20 | 122 |
| 3 | $\sim 105$ | 7 | 18 | 106 |
| 4 | $\sim 90$ | 6 | 15 | 91 |
| 5 | $\sim 75$ | 5 | 13 | 76 |
| 6 | $\sim 60$ | 4 | 10 | 61 |
| 7 | $\sim 45$ | 3 | 8 | 46 |
| 8 | $\sim 30$ | 2 | 5 | 30 |
| 9 | $\sim 15$ | 1 | 3 | 15 |
| 10 (Null) | 0 | 0 | 0 | 0 |

## S7 Shaping of chromatin structure by the spatial organization of CPCs

To complement the ensemble-averaged contact statistics, we quantified how CPCs distribute around the chromatin fiber and how this distribution relates to local folding. In Fig. S6, we measure CPC accessibility along the genome by the nearest-CPC distance. For each configuration, we define for bead *i*

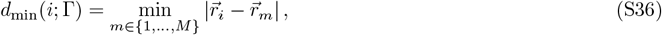

and report the ensemble average ⟨*d*_min_(*i*)⟩ as a function of genomic coordinate.

In the constant-connectivity setting with *N*_*c*_ ≃ 150 (Fig. S6a), low-valency systems contain many CPCs and therefore provide relatively uniform spatial access along the chain. As *k* increases, the number of CPCs decreases (Eq. (S14)), and the accessibility profile becomes more structured: beads partition into regions that remain consistently close to a CPC and regions that are farther away on average. Despite these differences, all valencies show the same qualitative trend that ⟨*d*_min_(*i*)⟩ anticorrelates with the local Hi-C activity (here taken as the Hi-C row-sum shown for reference). Thus, loci that contribute disproportionately to population-averaged contacts tend to be preferentially surrounded by CPCs, indicating that the inferred affinity landscape recruits CPCs to contact-rich genomic neighborhoods.

In fixed-abundance setting with *M* = 40 (Fig. S6b), high-valency systems show that each CPC can engage more loci, and accessibility remains broadly uniform across the region. At low valency, the limited total link capacity forces CPCs to concentrate on the strongest-affinity neighborhoods, producing pronounced peaks in ⟨*d*_min_(*i*)⟩ at loci that are comparatively disfavored by the inferred interaction blueprint. This spatial inhomogeneity provides a direct structural signature of a link-capacity-limited regime and is consistent with the degraded Hi-C agreement observed for low *k* at fixed *M* (Fig. S4).

Finally, Fig.S6c,d illustrate how fluctuations in CPC localization couple to local structural “breathing”. By grouping conformations according to where CPCs are most enriched along the polymer (Fig.S6c) and comparing the cluster-averaged distance patterns to the global average (Fig. S6d), we find a direct correspondence between CPC crowding and local compaction. When CPCs transiently accumulate in a given subTAD, that region becomes locally more compact, reflected by reduced spatial distances (and increased local contact density) relative to the ensemble mean. Conversely, conformations with a more dispersed CPC distribution show weaker local compaction. These snapshots indicate that CPCs do not impose purely static constraints. Instead, their stochastic redistribution along the polymer provides a natural mechanism for time-dependent, region-specific condensation, contributing to the observed single-cell variability even under the same population-averaged constraints. This coupling becomes more pronounced as CPC valency increases, consistent with high-valency CPCs promoting more intermittent clustering and stronger domain-level breathing.

## S8 Hybrid simulations: mixed-valency CPC pools

To examine whether chromatin folding benefits from combining low- and high-valency CPCs, we constructed hybrid CPC-MEM systems that contain both *CPC*^2^ and *CPC*^6^. Across all hybrid compositions, we kept the connectivity budget approximately fixed at *N*_*c*_ ≃ 150 by adjusting CPC counts according to

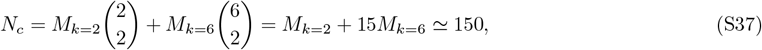

where *M*_2_ and *M*_6_ denote the numbers of *CPC*^2^ and *CPC*^6^, respectively (Table S2). Starting from the pure-*CPC*^2^ system, we progressively replaced *CPC*^2^ with *CPC*^6^ while maintaining the same total link capacity.

**Table S2.** Composition of hybrid systems. The numbers of high-valency (*M*_*k*=6_) and low-valency (*M*_*k*=2_) CPCs are adjusted to maintain a nearly constant Connectivity budge (*N*_*c*_ ≈ 150).

| Mixture ID | $M_{k=6}$ | $M_{k=2}$ | Total $N_c$ |
| --- | --- | --- | --- |
| Pure $k = 2$ | 0 | 152 | 152 |
| Mix 1 | 1 | 137 | 152 |
| Mix 2 | 2 | 122 | 152 |
| Mix 3 | 3 | 107 | 152 |
| Mix 4 | 4 | 92 | 152 |
| Mix 5 | 5 | 77 | 152 |
| Mix 6 | 6 | 62 | 152 |
| Mix 7 | 7 | 47 | 152 |
| Mix 8 | 8 | 32 | 152 |
| Mix 9 | 9 | 17 | 152 |
| Pure $k = 6$ | 10 | 0 | 150 |

Each mixture composition is treated as an independent maximum-entropy inference problem at fixed connectivity budget *N*_*c*_ ≃ 150. Along the titration series we replace a subset of *CPC*^2^ by *CPC*^6^ while adjusting bead counts so that the maximum link capacity 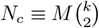 remains approximately constant across mixtures (Table S2). In contrast to depletion simulations, where the learned landscape is held fixed, we re-optimize the affinity matrix {*α*_*ij*_} for every hybrid system against the same experimental Hi-C target, so that each ensemble represents the equilibrium structure implied by that specific mixed-valency pool under the maximum-entropy constraint.

Fig. S7 summarizes how population-averaged and single-cell metrics evolve with mixture composition. For each hybrid, we report the ensemble-averaged contact probability map *P*_*ij*_ and mean spatial-distance matrix ⟨*r*_*ij*_⟩ (Fig. S7a,b), the Hi-C reconstruction error as a function of mixing ratio (Fig. S8c), and standard validation/readout metrics including *P*(*s*) (Fig. S7d), probe-pair distance distributions (Fig. S7e), the separation-score analysis (Fig. S8f), and distributions of *n*_*c*_, dRMSD, and *V*_eff_ (Fig. S8g–i). Across the titration series, enriching the mixture with *CPC*^6^ systematically improves map-level agreement, while shifting the ensemble toward greater structural variability, reflecting the reduced constraint efficiency per pairwise link at higher valency under fixed *N*_*c*_.

Fig. S8 visualizes how the mixed pool realizes this trade-off at the level of contact topology and CPC organization. Composite maps along the titration series show that increasing the *CPC*^6^ fraction introduces stronger, more localized co-bridging features, whereas the *CPC*^2^ population contributes a dense background of pairwise bridges that supports globally cohesive packing (Fig. S8a). We further characterize a representative intermediate mixture 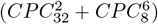 by reporting an effective connected-valency readout (Fig. S8b), loop-span distributions decomposed by CPC type (Fig. S8c), and radial density profiles of CPCs and chromatin relative to the chromatin center of mass (Fig. S8d). Together, Figs. S7 and S8 show that mixed valencies provide a direct route to balance population-level contact fidelity and single-cell structural variability under a fixed connectivity budget.

## S9 Perturbation simulations: removing boundary-associated affinities

To assess how boundary and anchor features encoded in the inferred affinity landscape contribute to 3D chromatin organization, we performed targeted perturbation simulations of the optimized parameters. We identified a set of loci with strong CTCF boundary/anchor signal from Fig. 3h and defined the corresponding bead-index set

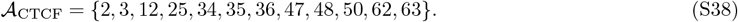

Rather than altering the polymer representation, we constructed a “mutant” affinity matrix by removing interactions involving these CTCF-enriched bins from the learned landscape:

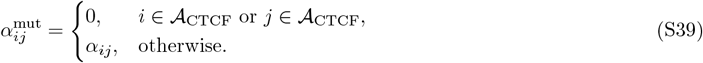

We applied the same procedure to the optimized {*α*_*ij*_} obtained for the *CPC*^2^, *CPC*^4^, *CPC*^6^, and mixed systems. The mixed system corresponds to a heterogeneous CPC pool containing 32 *CPC*^2^ and 8 *CPC*^6^. For each case, we reran simulations using 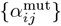 without re-optimizing parameters, so that any structural changes reflect the loss of boundary-associated affinities under an otherwise fixed interaction blueprint.

Fig. S9 summarizes the structural consequences. The differential contact maps (Fig. S9a, lower triangles) show a localized loss of contacts concentrated around the perturbed boundary and anchor indices, confirming that interactions involving 𝒜_CTCF_ contribute disproportionately to stabilizing the corresponding architectural features. Consistent with this contact loss, the locus-resolved distance patterns relax in a nonuniform manner (Fig. S9b): distances across the affected boundary and anchor neighborhoods increase most strongly, whereas short-range intra-domain distances are comparatively less perturbed, indicating that the perturbation primarily weakens boundary-level organization rather than uniformly decompacting the chain.

Global compaction changes are modest at the level of overall size, but local packing and connectivity are reduced. This is reflected by a downward shift in the contacts-per-bead distribution *n*_*c*_ (Fig. S9f), which reports a net loss of realized CPC-mediated links in the mutant ensembles. Importantly, the insulation responds in the expected direction: boundary probability profiles show weakened boundary signatures upon perturbation (Fig. S9e). The effect is most pronounced for the low-valency *CPC*^2^ system, consistent with the fact that distributed pairwise bridging relies sensitively on the learned boundary-linked affinities, whereas high-valency, hub-dominated systems retain a larger fraction of their connectivity through multi-way clustering outside the perturbed set.

Overall, selectively zeroing interactions associated with strong CTCF boundary and anchor loci attenuates boundary insulation and locally relaxes the corresponding loop- and domain-scale organization, while leaving much of the remaining intra-domain packing comparatively intact. This supports the interpretation that {*α*_*ij*_} encodes locus-specific architectural “blueprint” information, and that boundary and anchor-linked entries constitute a key subset of constraints that stabilize the corresponding TAD-scale features.

## S10 Dissecting heterogeneity: control with a uniform affinity landscape

To disentangle the role of locus specificity from the role of CPC constraint topology, we constructed a control in which the optimized affinity landscape {*α*_*ij*_} was replaced by a non-specific baseline that depends only on genomic separation. Concretely, we set *α*_*ij*_ → *α*(|*i* − *j*|), thereby removing locus identity while preserving the overall separation-dependent trend (Fig. S10a). We then applied the same sampling and analysis pipeline as for the locus-specific models and compared ensembles across valencies.

As expected, the uniform-affinity control retains coarse polymeric scaling and reproduces the global distance-dependent decay of contacts (Fig. S10b,c,e), but it cannot encode architectural features that require positional information. The resulting contact maps therefore lose locus-resolved structure: loop- and domain-like features are strongly attenuated, and boundary statistics become largely featureless (Fig. S10g). The same limitation appears in the imaging validation. Probe-pair distance distributions that are selectively constrained in the optimized model broaden and shift under uniform *α*, because the energetic preference stabilizing those contacts is replaced by a generic separation-only interaction (Fig. S10h).

The heterogeneity analysis clarifies what is intrinsic to valency and what requires a locus-specific blueprint. Across the uniform-*α* ensembles, higher valency still produces larger conformational variability than lower valency (Fig. S10i), indicating that valency-controlled constraint topology contributes a landscape-independent component to structural heterogeneity. In contrast, the absolute dRMSD levels increase for all valencies and exceed the experimental benchmark (Fig. S10j), showing that the locus-specific {*α*_*ij*_} acts as an informational constraint that confines the conformational manifold and suppresses excess entropy. Together, these results emphasize that realistic chromatin ensembles require two ingredients: a structured, Hi-C-inferred affinity landscape that specifies where interactions are favored, and CPC-mediated constraint topology (set by valency and abundance) that determines how efficiently those preferences are assembled into geometric constraints and residual plasticity.

## S11 2D demonstrative model for valency effects

To build intuition for how interaction valency reshapes chromatin folding, we implemented a minimal two-dimensional (2D) polymer-binder model with explicit, transient binders. This toy model is not used for inference; it is only meant to visualize, in real time, how changing valency alters the topology of constraints and the resulting polymer conformational variability.

Unlike CPC-MEM, which samples equilibrium ensembles using Metropolis–Hastings Monte Carlo, the 2D model is simulated by Langevin dynamics. The position 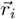 of each polymer bead and binder bead evolves as

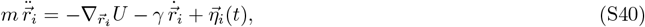

where *γ* is a friction coefficient and 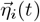 is Gaussian thermal noise with zero mean and variance chosen to satisfy fluctuation-dissipation at the target temperature.

The chromatin fiber is represented as a bead-spring chain in 2D. Neighboring beads are connected by harmonic springs, and all polymer beads and binders interact through a soft repulsive potential to prevent overlaps. Binders diffuse freely and can form transient harmonic tethers to nearby chromatin beads through a capture-and-release mechanism. Specifically, a bond between a binder and a bead is created when their distance falls below *R*_bind_, provided the binder has not yet reached its valency limit. This rule strictly enforces that each binder carries at most *k* simultaneous bonds at any time. To mimic finite interaction lifetimes, each existing bond is removed stochastically with a probability *P*_off_ per integration step.

To parallel the comparisons with the 3D MC simulation results, we simulated two distinct conditions. In the first scenario, we maintained a constant connectivity budget (*N*_*c*_ = 90; Fig. S11). Under this constraint, bivalent binders (*k* = 2) generate relatively uniform compaction, whereas higher valency binders (*k* = 4, 6) concentrate links into hubs and produce more heterogeneous conformations. In the second scenario, we fixed the total binder number (*M* = 10; Fig. S12). Here, the bivalent system becomes connectivity-limited and cannot efficiently compact the polymer, while increasing *k* raises the available link capacity and restores compaction.

Together, the 2D model provides a direct visual counterpart of the core mechanism emphasized by CPC-MEM: at fixed connectivity, increasing valency redistributes links into locally redundant hubs and increases structural variability, whereas at fixed binder abundance, increasing valency primarily increases link capacity and strengthens compaction.

**Figure S1.**
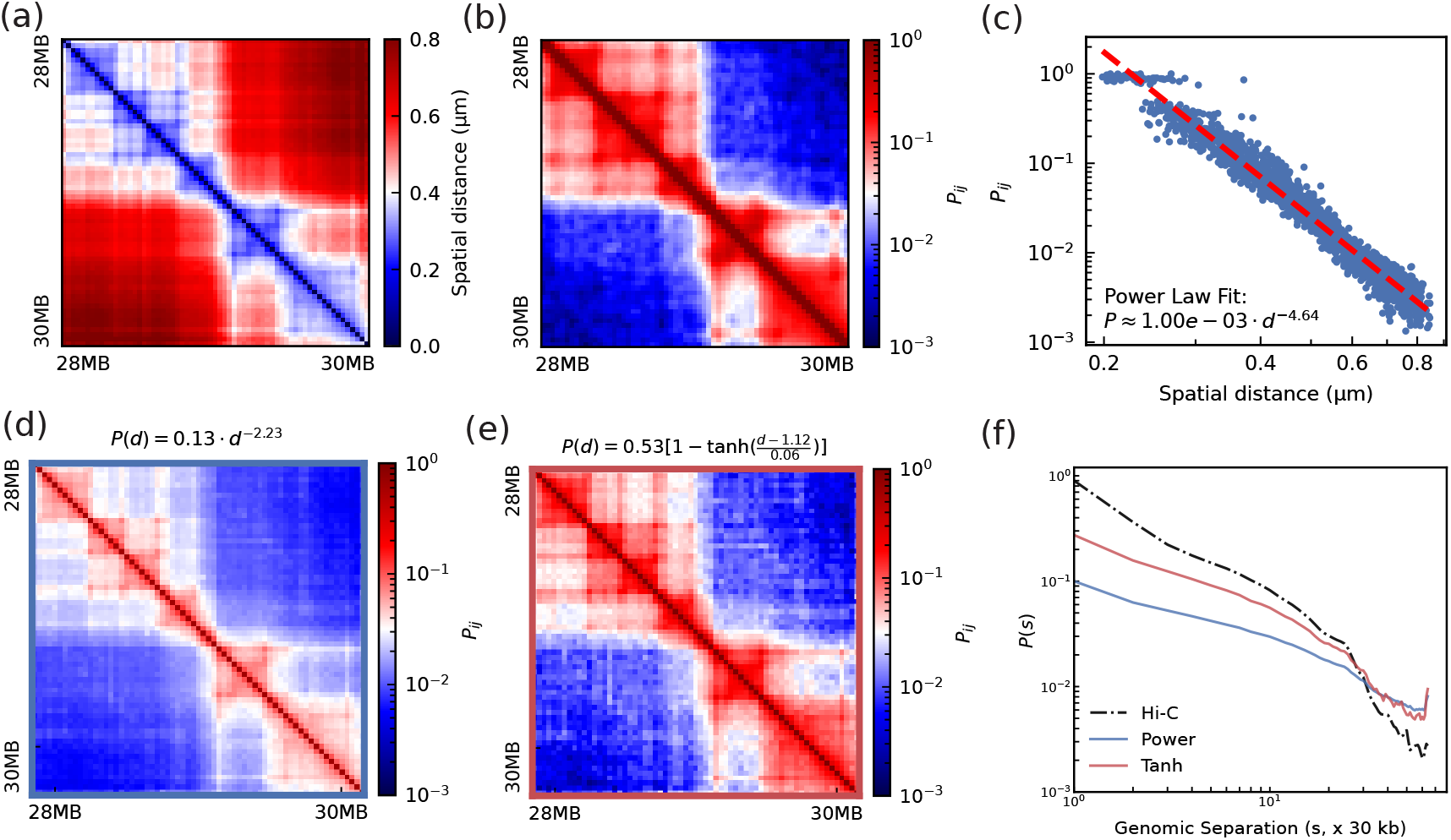
Calibration of the distance-contact mapping between Hi-C and imaging. (a) Hi-C contact-probability matrix for the studied region. (b) Mean spatial-distance matrix computed from super-resolution chromatin tracing. (c) Relationship between contact probability and mean spatial distance for all locus pairs; the red dashed line shows a power-law fit. (d) Contact map reconstructed from imaging coordinates using a power-law mapping. (e) Same as (d), but using a tanh-based mapping that better captures the full dynamic range. (f) Genomic-distance decay *P*(*s*) for the experimental Hi-C map and the reconstructed maps in (d,e), highlighting the improved agreement for the tanh mapping.

**Figure S2.**
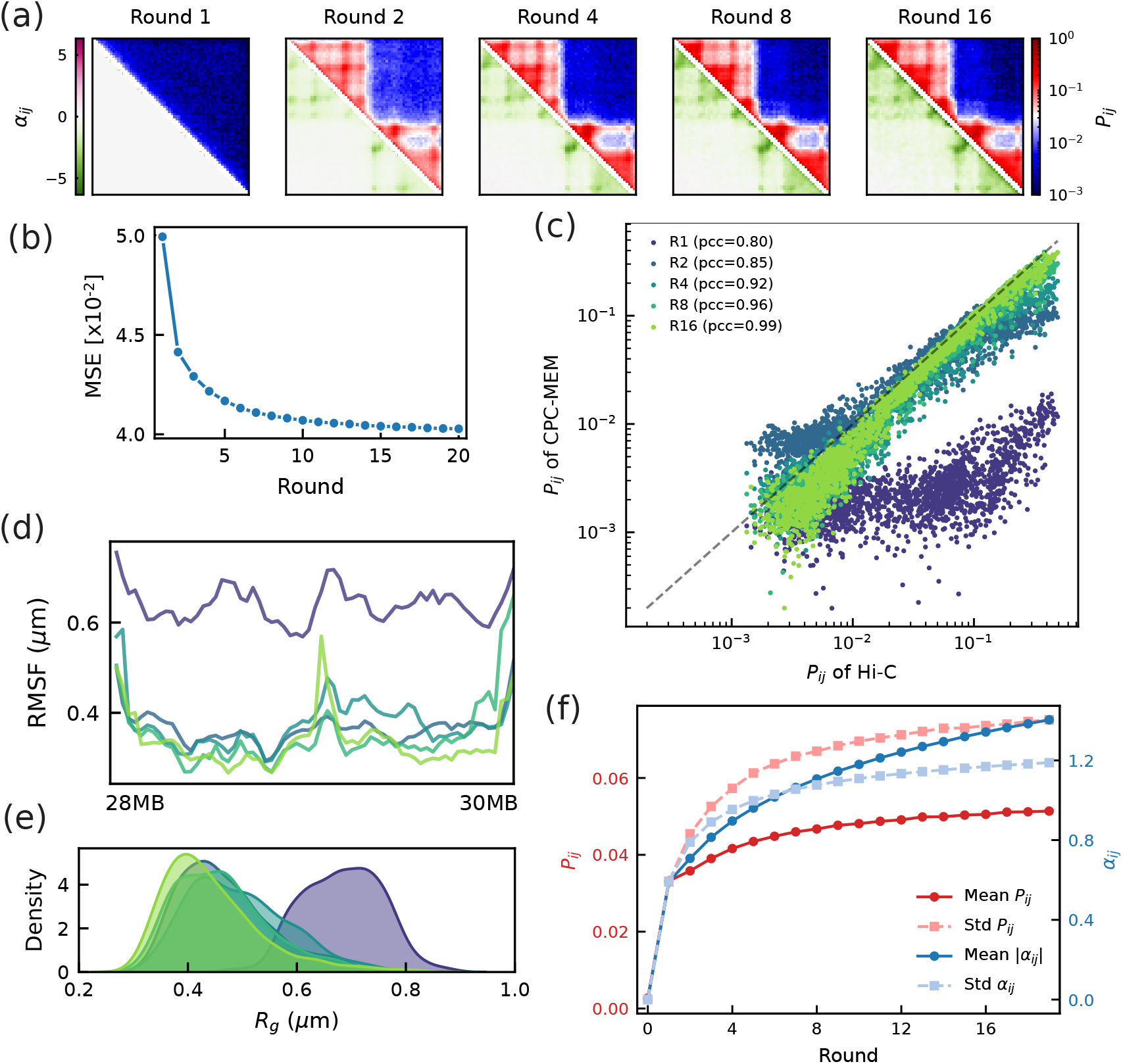
Iterative maximum-entropy optimization for a representative CPC-MEM system 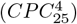. (a) Evolution across optimization rounds of the inferred affinity matrix {*α*_*ij*_} (upper triangle) and the corresponding simulated contact-probability map (lower triangle), starting from {*α*_*ij*_} = 0. (b) Training loss (MSE between simulated and experimental Hi-C contact probabilities) as a function of round. (c) Round-by-round agreement between simulated and experimental contact probabilities shown as scatter plots (log scale). (d) Bead-resolved root-mean-square fluctuation (RMSF) profiles reporting changes in positional mobility during optimization. (e) Distributions of the radius of gyration *R*_*g*_ across rounds, illustrating the accompanying global compaction. (f) Convergence assessment showing the evolution of the mean and standard deviation of contact probabilities (red, left axis) and of inferred parameters {*α*_*ij*_} = 0 (blue, right axis).

**Figure S3.**
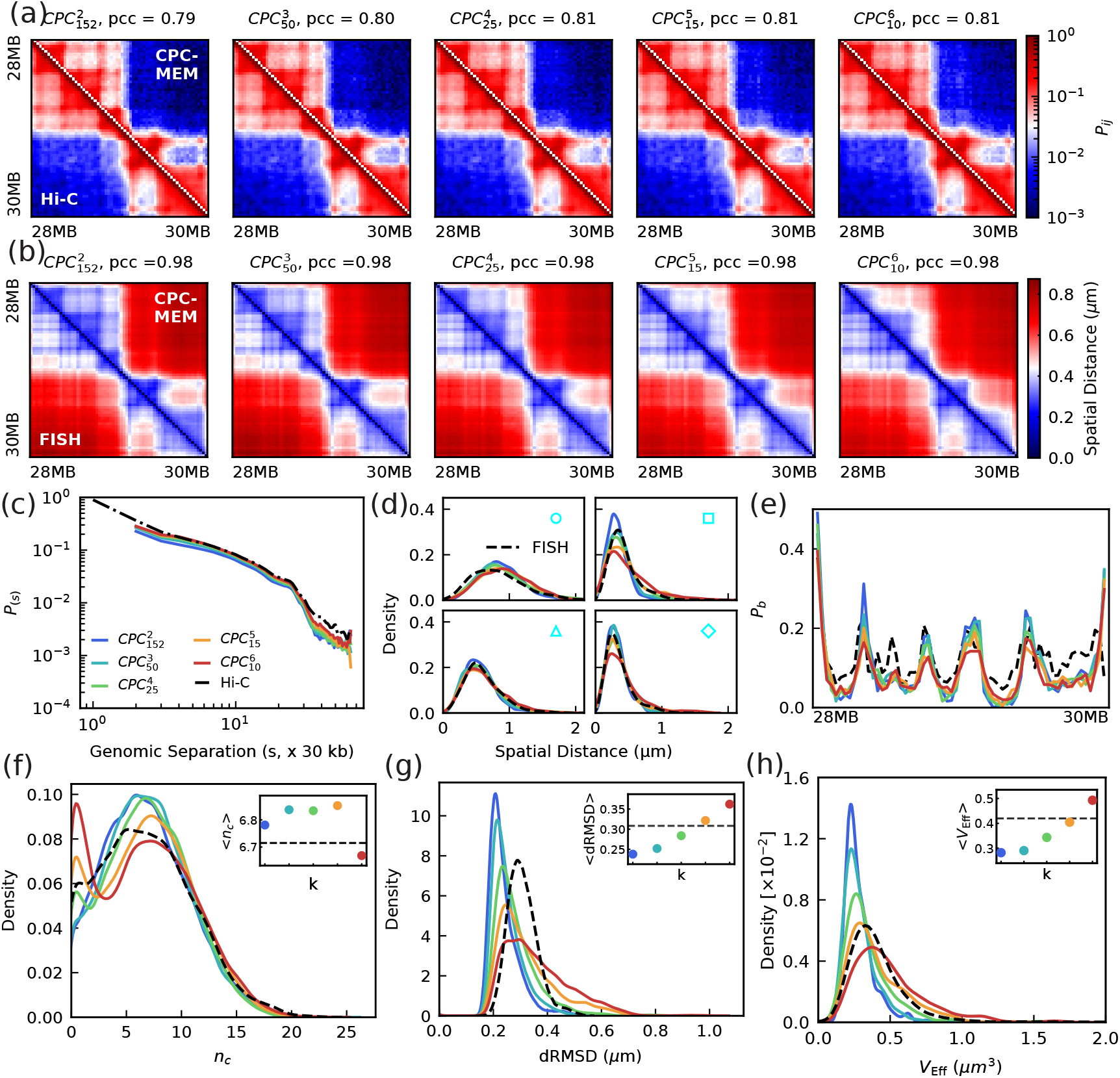
Valency dependence of CPC-MEM ensembles at fixed connectivity budget (*N*_*c*_ ≃ 150). (a) Ensemble-averaged contact probability maps *P*_*ij*_ inferred for CPC valencies *k* = 2, …, 6 (the same experimental Hi-C map is used as the training target for all cases; *M* is adjusted with *k* to keep *N*_*c*_ constant). Agreement with Hi-C is shown as Pearson correlation coefficient (*pcc*) reported above each map. (b) Corresponding mean spatial-distance matrices ⟨*r*_*ij*_⟩ from the simulated ensembles. (c) Genomic-separation dependence of contact probability, *P*(*s*). (d) Distributions of inter-locus distances for the probe pairs defined in Fig. 1(b), compared with imaging. (e) Boundary probability profiles *P*_*b*_(*i*) extracted from single-snapshot separation-score analysis. (f–h) Distributions of per-locus link number *n*_*c*_(*i*), structural heterogeneity quantified by dRMSD, and effective volume *V*_eff_. Insets report the corresponding mean values as a function of *k*; dashed lines mark experimental references where available.

**Figure S4.**
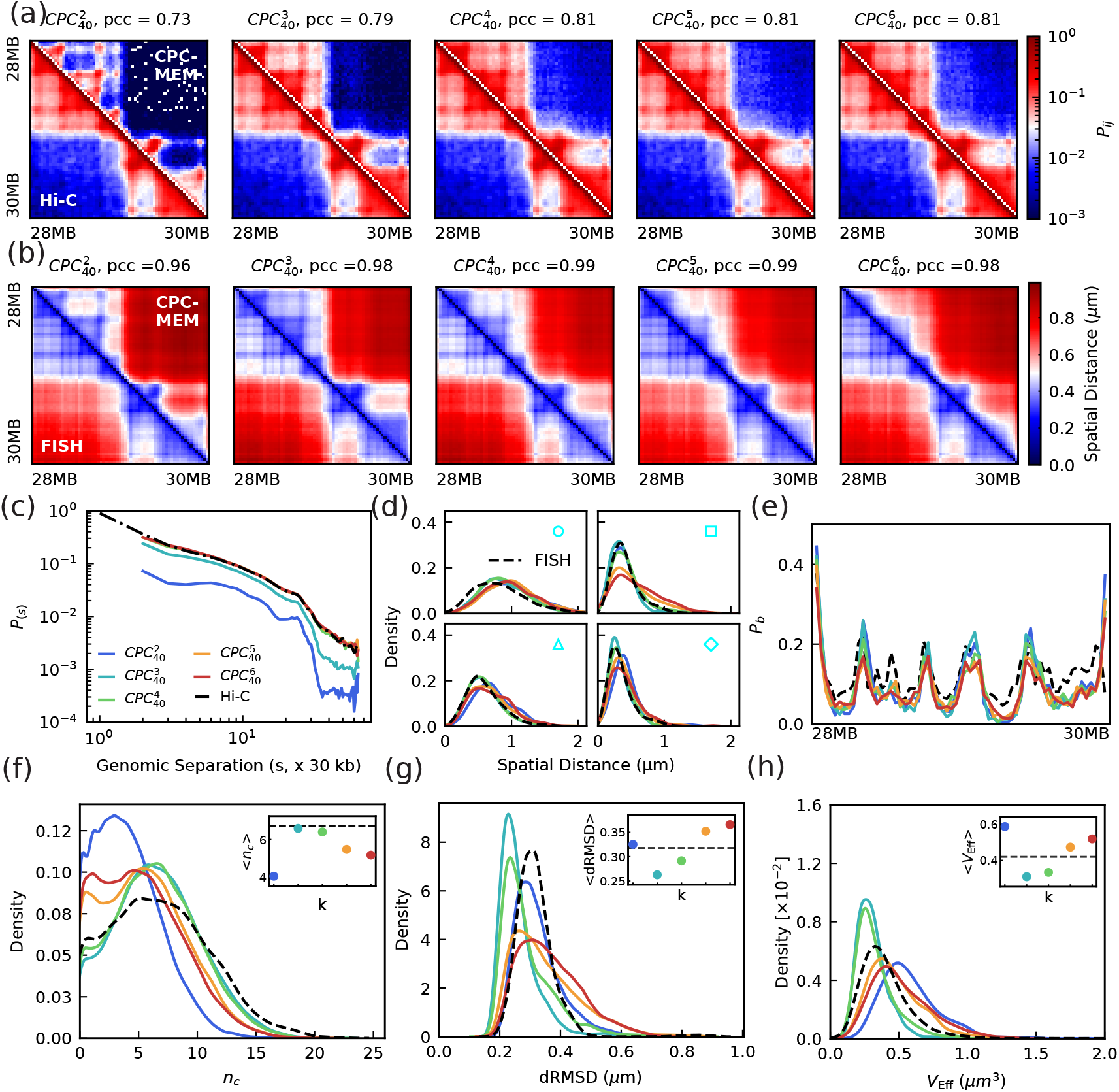
Structural properties of CPC-MEM ensembles at fixed CPC abundance (*M* = 40). With CPC number held constant, the theoretical connectivity budget increases with valency as 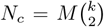 (from *N*_*c*_ = 40 at *k* = 2 to *N*_*c*_ = 600 at *k* = 6). The analysis and panel layout follow Fig. S3. (a,b) Ensemble-averaged contact probability maps *P*_*ij*_ and mean spatial-distance matrices ⟨*r*_*ij*_⟩ for *k* = 2–6 (correlations with Hi-C and imaging shown above). (c) Genomic-separation dependence of contact probability, *P*(*s*). (d) Distributions of inter-locus distances for the probe pairs defined in Fig. 1(b), compared with imaging. (e) Boundary probability profiles *P*_*b*_(*i*) extracted from single-snapshot separation-score analysis. (f–h) Distributions of per-locus link number *n*_*c*_(*i*), structural heterogeneity quantified by dRMSD, and effective volume *V*_eff_. Insets report the corresponding mean values as a function of *k*; dashed lines mark experimental references where available.

**Figure S5.**
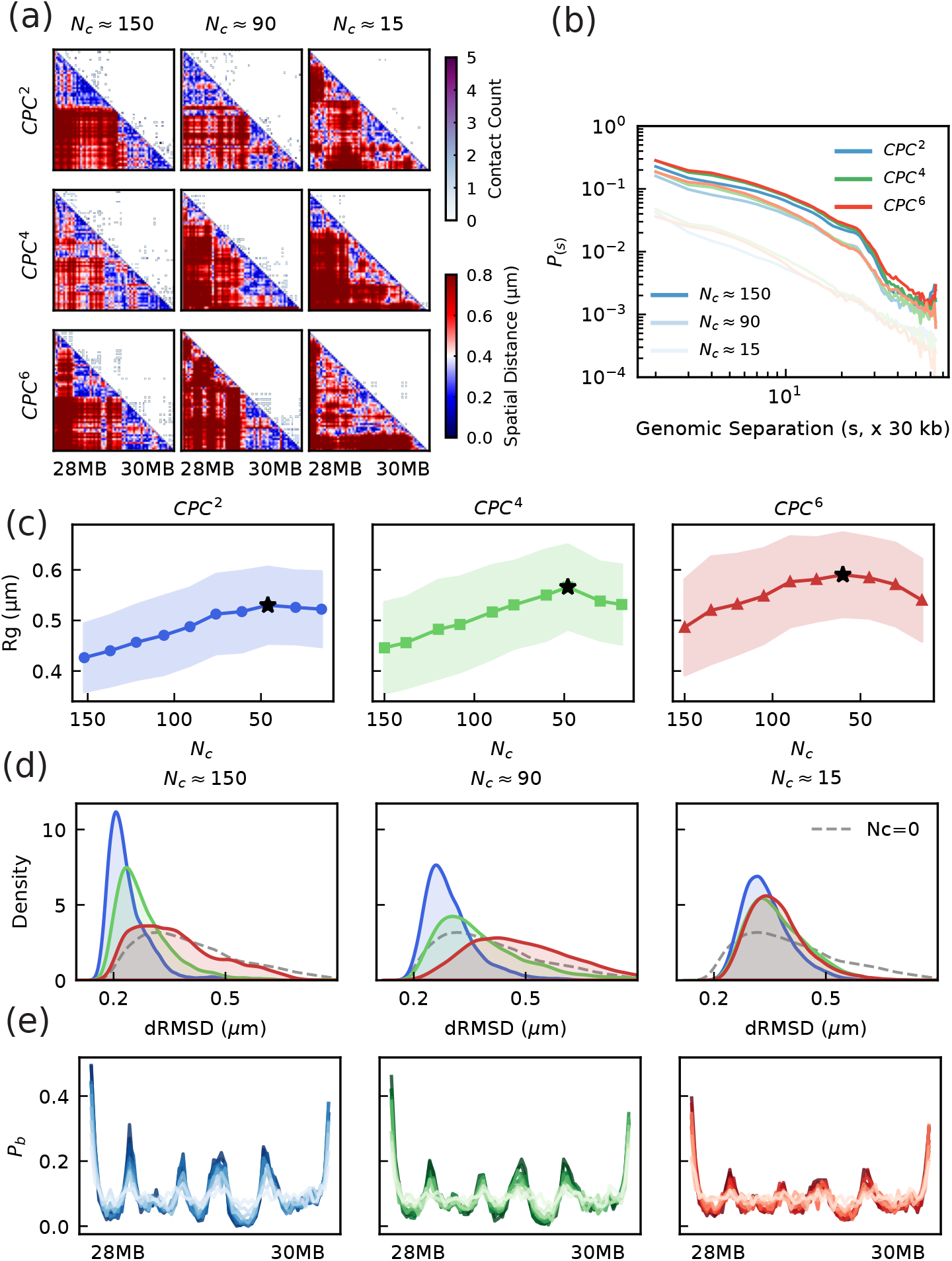
Structural evolution under CPC depletion at fixed learned affinity landscape. CPC-MEM models of different valency are first optimized at the saturated reference state (*N*_*c*_ ≃ 150) and then titrated by reducing CPC abundance while keeping {*α*_*ij*_} fixed. (a) For each valency and titration point, matrices show CPC-mediated contact counts (upper triangle) and the corresponding mean spatial distances (lower triangle). (b) Contact-probability decay *P*(*s*) during depletion, highlighting the progressive loss of long-range contacts at decreasing *N*_*c*_. (c) Mean radius of gyration *R*_*g*_ versus *N*_*c*_; shaded bands indicate the standard deviation across the ensemble and reveal substantially larger conformational fluctuations for high valency compared with the low-valency scaffold. (d) dRMSD distributions quantifying ensemble heterogeneity; the pronounced broadening and heavy tail at high valency indicate increased structural plasticity under the same connectivity loss. (e) Boundary probability profiles *P*_*b*_(*i*) across depletion; color intensity (dark to light) denotes decreasing *N*_*c*_, illustrating the erosion of boundary recurrence as CPC-mediated links are removed.

**Figure S6.**
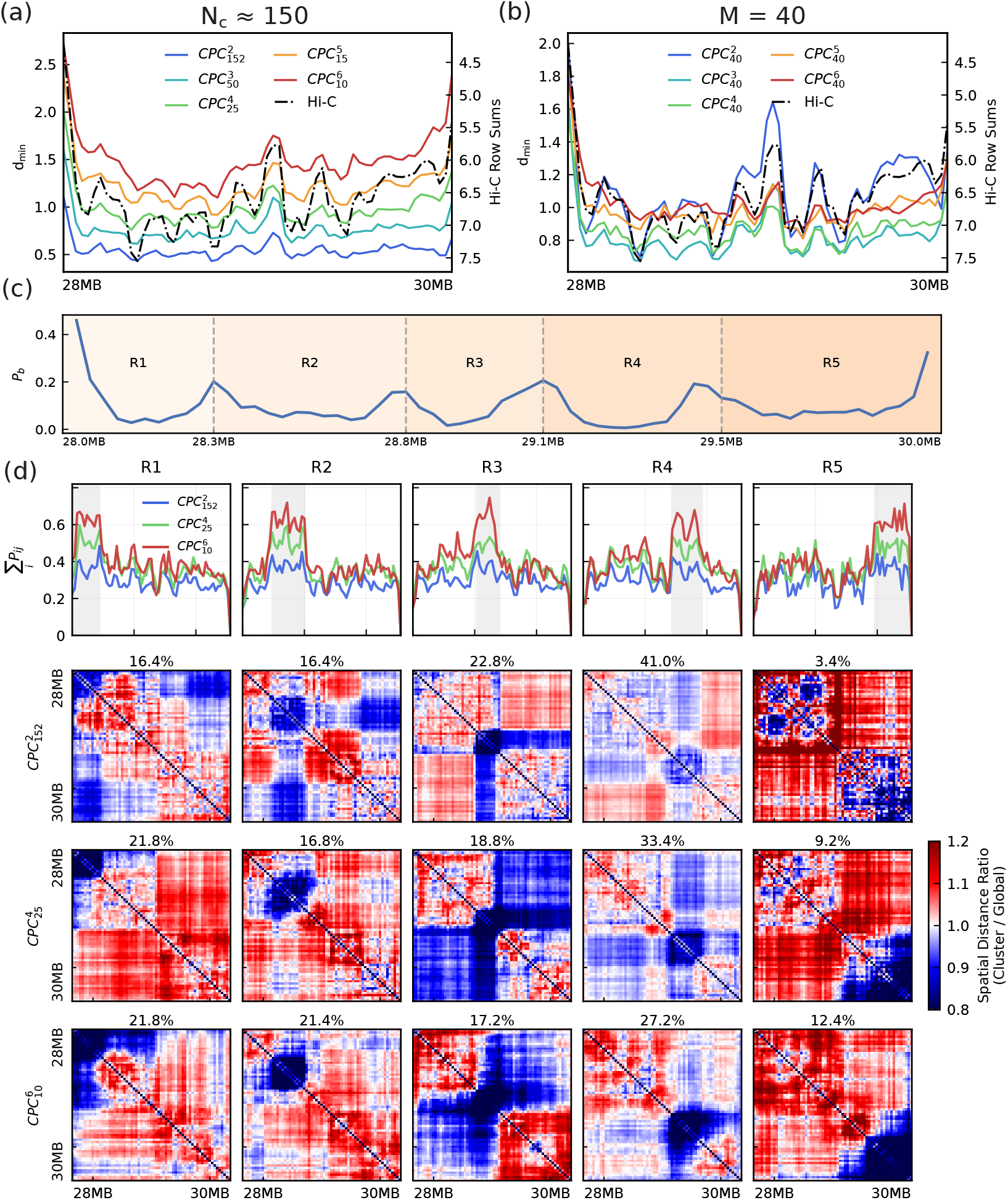
Spatial organization of CPCs and its coupling to local chromatin compaction. (a) Nearest-CPC accessibility along the genome in the constant-connectivity comparison (*N*_*c*_ ≃ 150), quantified by the ensemble-averaged distance from each bead to its closest CPC. The dashed curve shows the (inverted) experimental Hi-C row sum for the same region as a 1D proxy for loci with high contact participation. (b) Same analysis as in (a) for the constant-abundance comparison (*M* = 40). (c) Boundary probability profile and subTAD partitions used to assign conformations by the location of maximal contact density. (d) Cluster-resolved organization for the constant-connectivity ensembles. Conformations are grouped by the subTAD carrying the maximal contact density (using the partitions in (c)). Top: cluster-averaged contact-sum profiles. Bottom: ratios of mean spatial-distance matrices, ⟨*d*_*ij*_⟩_cluster_/ ⟨*d*_*ij*_⟩_global_, where values < 1 indicate local compaction and values > 1 indicate local expansion relative to the full ensemble. Percentages in the subtitles report the cluster weights.

**Figure S7.**
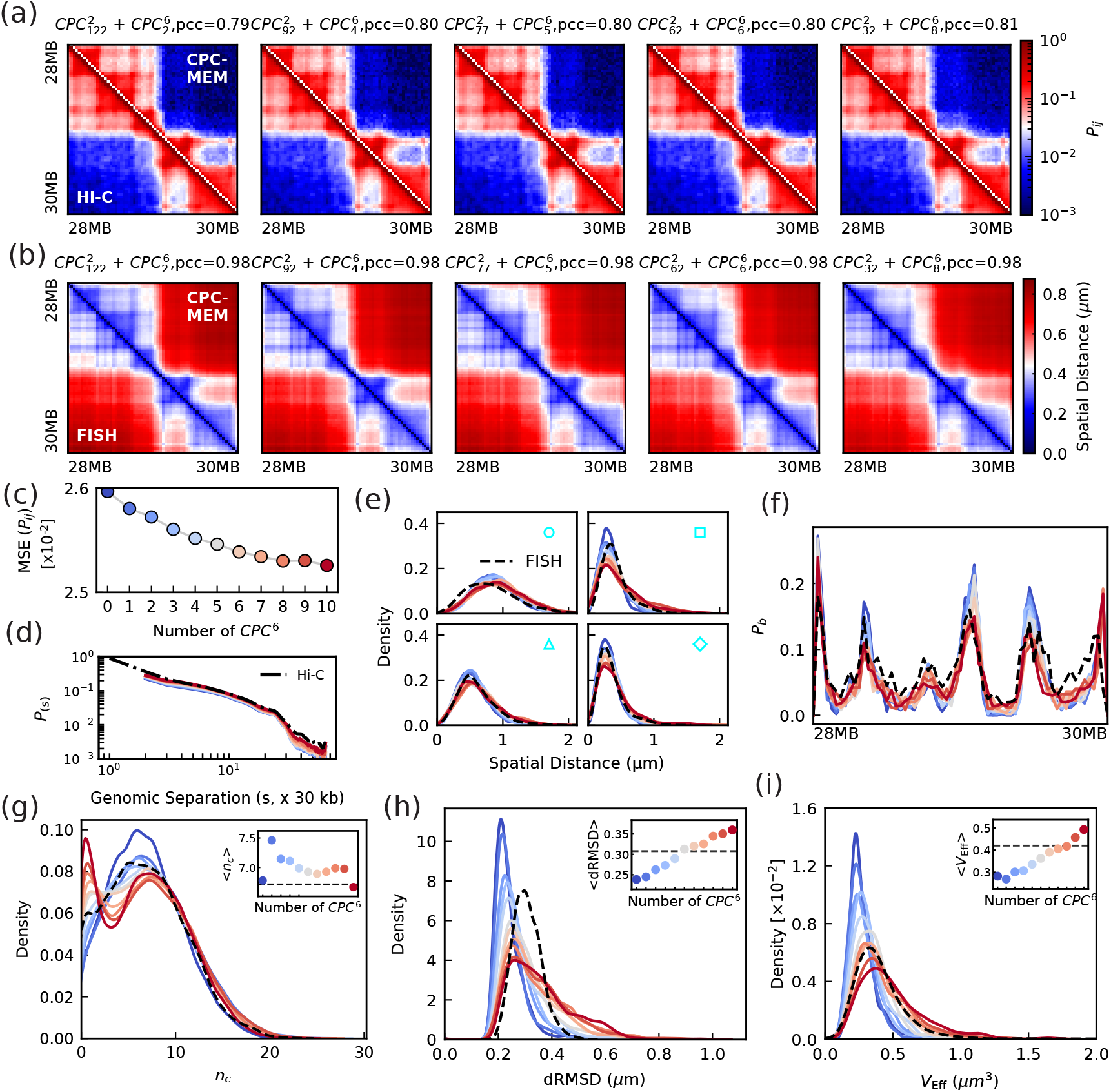
Structural properties of mixed-valency CPC-MEM ensembles across mixture compositions. The analysis and panel layout follow Fig. S3. (a,b) Ensemble-averaged contact probability maps *P*_*ij*_ and mean spatial-distance matrices ⟨*r*_*ij*_⟩; correlations to the experimental Hi-C and imaging references are reported above each column. (c) Mean-squared error (MSE) between simulated and experimental *P*_*ij*_ as a function of the mixing ratio. (d) Genomic-separation dependence of contact probability, *P*(*s*). (e) Distributions of inter-locus distances for the probe pairs defined in Fig. 1(b), compared with imaging. (f) Boundary probability profiles *P*_*b*_(*i*) extracted from single-snapshot separation-score analysis. (g–i) Distributions of per-locus link number *n*_*c*_(*i*), structural heterogeneity quantified by dRMSD, and effective volume *V*_eff_. Insets report the corresponding mean values as a function of number of *CPC*^6^; dashed lines mark experimental references where available.

**Figure S8.**
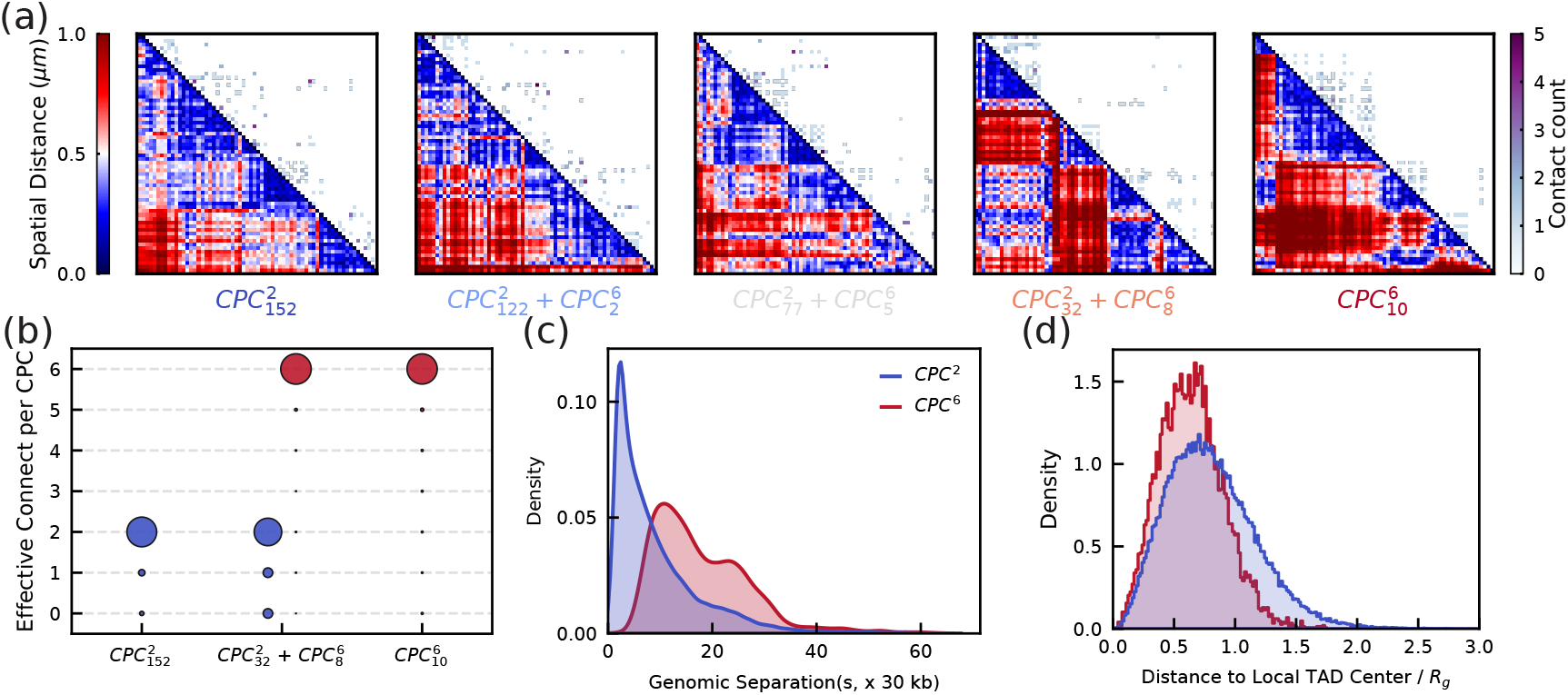
Mixed-valency CPC ensembles at fixed connectivity budget (*N*_*c*_ ≃ 150). (a) Representative composite maps for mixtures interpolating from pure *CPC*^2^ to pure *CPC*^6^ while keeping *N*_*c*_ approximately constant by adjusting CPC counts. Upper triangles show contact account maps; lower triangles show spatial-distance matrices. (b–d) Additional diagnostics for a representative hybrid mixture 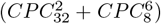. Shown are the effective connected valency (b), loop-span distributions decomposed by CPC type (c), and radial density profiles of CPCs and chromatin measured relative to the chromatin center of mass (d).

**Figure S9.**
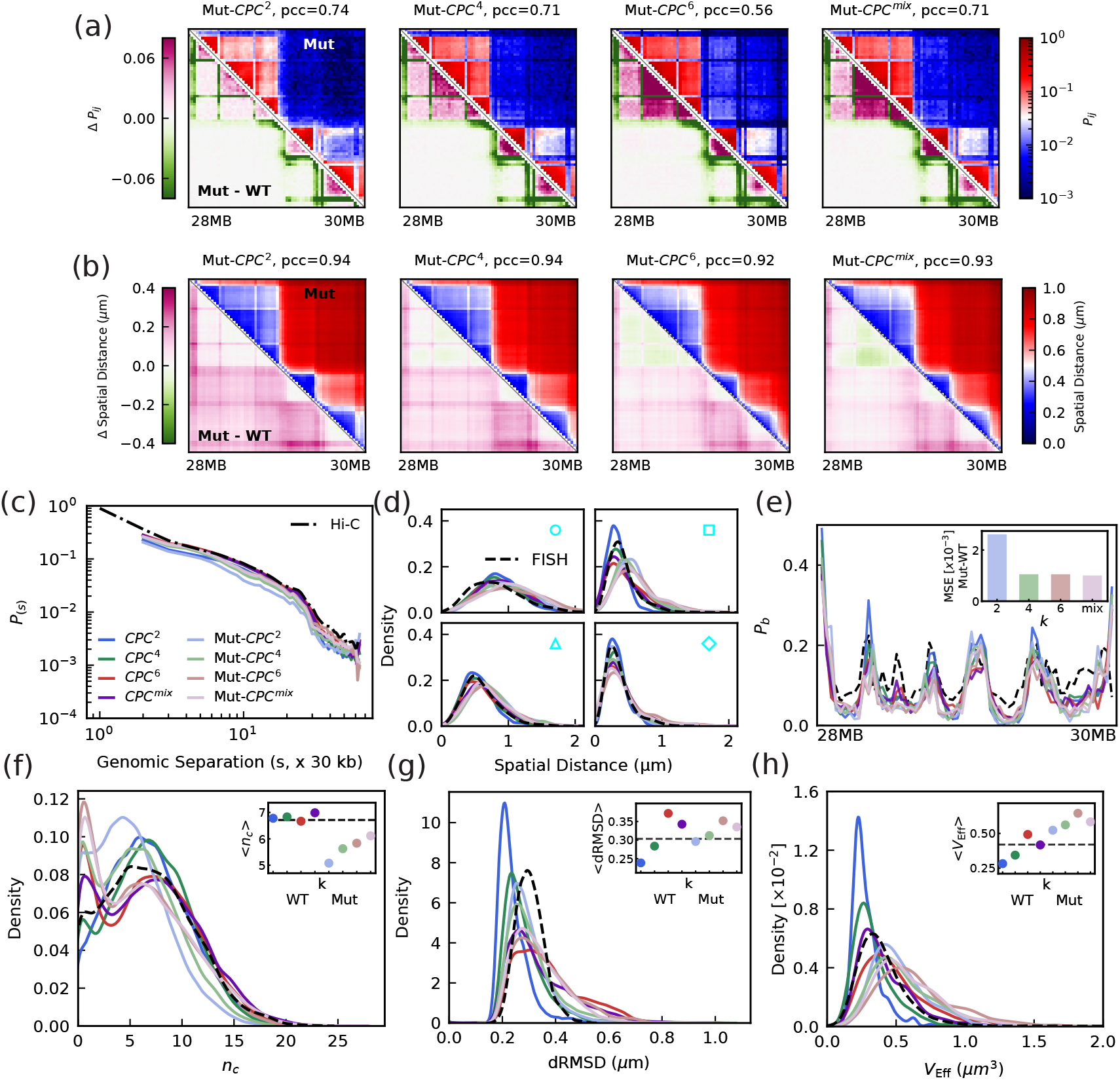
Effects of removing CTCF-associated interactions from the learned affinity landscape. Mutant simulations were performed using a perturbed parameter matrix 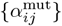 in which all interactions involving CTCF-enriched boundary/anchor bins were removed (bead indices *B* = {2, 3, 12, 25, 34, 35, 36, 47, 48, 50, 62, 63}); all other entries were unchanged. The analysis and panel layout follow Fig. S3. (a) Ensemble-averaged contact-probability maps *P*_*ij*_. (b) Mean spatial-distance matrices ⟨*r*_*ij*_⟩. In (a,b), upper triangles show the mutant ensemble averages, and lower triangles show differences relative to the wild type (Mutant WT). (c) Genomic-separation dependence of contact probability, *P*(*s*). (d) Distributions of inter-locus distances for the probe pairs defined in Fig. 1(b), compared with imaging. (e) Boundary probability profiles *P*_*b*_(*i*) extracted from single-snapshot separation-score analysis; inset reports the MSE between mutant and WT profiles. (f–h) Distributions of per-locus link number *n*_*c*_(*i*), structural heterogeneity quantified by dRMSD, and effective volume *V*_eff_. Insets report the corresponding mean values as a function of *k*; dashed lines mark experimental references where available.

**Figure S10.**
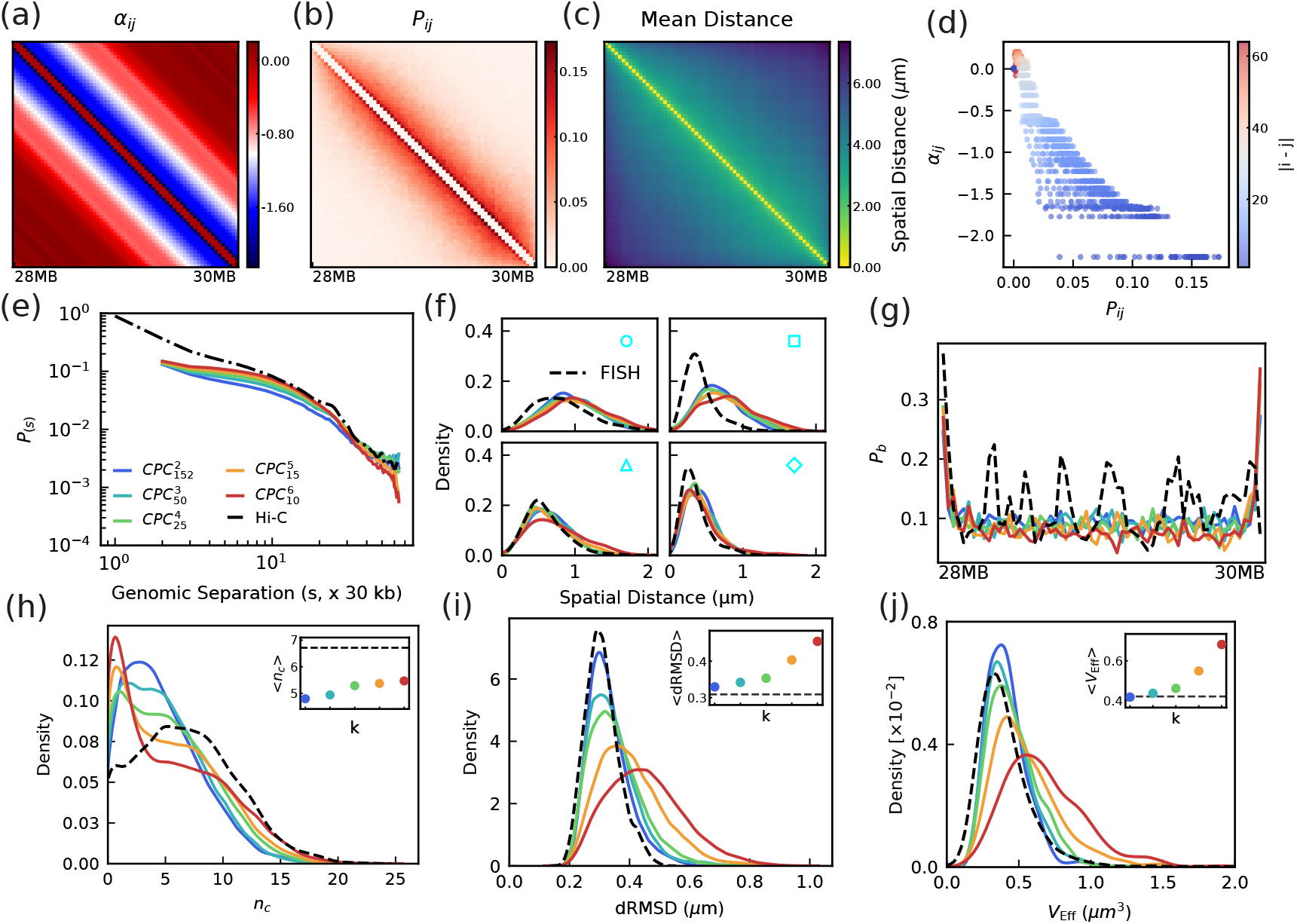
Uniform-affinity control (Uniform-*α*) decouples valency-driven constraint topology from locus-specific interaction patterns. The optimized, locus-resolved affinity landscape is replaced by a separation-only baseline *α*_*ij*_ = *α*(|*i* − *j*|), while all other model settings and analysis metrics are kept identical to the locus-specific CPC-MEM runs. The analysis and panel layout follow Fig. S3. (a) Input baseline affinity matrix {*α*_*ij*_}. (b,c) Ensemble-averaged contact-probability map *P*_*ij*_ and mean spatial-distance matrix ⟨*r*_*ij*_⟩. (d) Scatter plot of inferred contacts versus affinity (*P*_*ij*_ vs. *α*_*ij*_) under the uniform landscape. (e) Genomic-separation dependence of contact probability, *P*(*s*). (f) Distributions of inter-locus distances for the probe pairs defined in Fig. 1(b), compared with imaging. (g) Boundary probability profiles *P*_*b*_(*i*) extracted from single-snapshot separation-score analysis. (h–j) Distributions of per-locus link number *n*_*c*_(*i*), structural heterogeneity quantified by dRMSD, and effective volume *V*_eff_. Insets report the corresponding mean values as a function of *k*; dashed lines mark experimental references where available.

**Figure S11.**
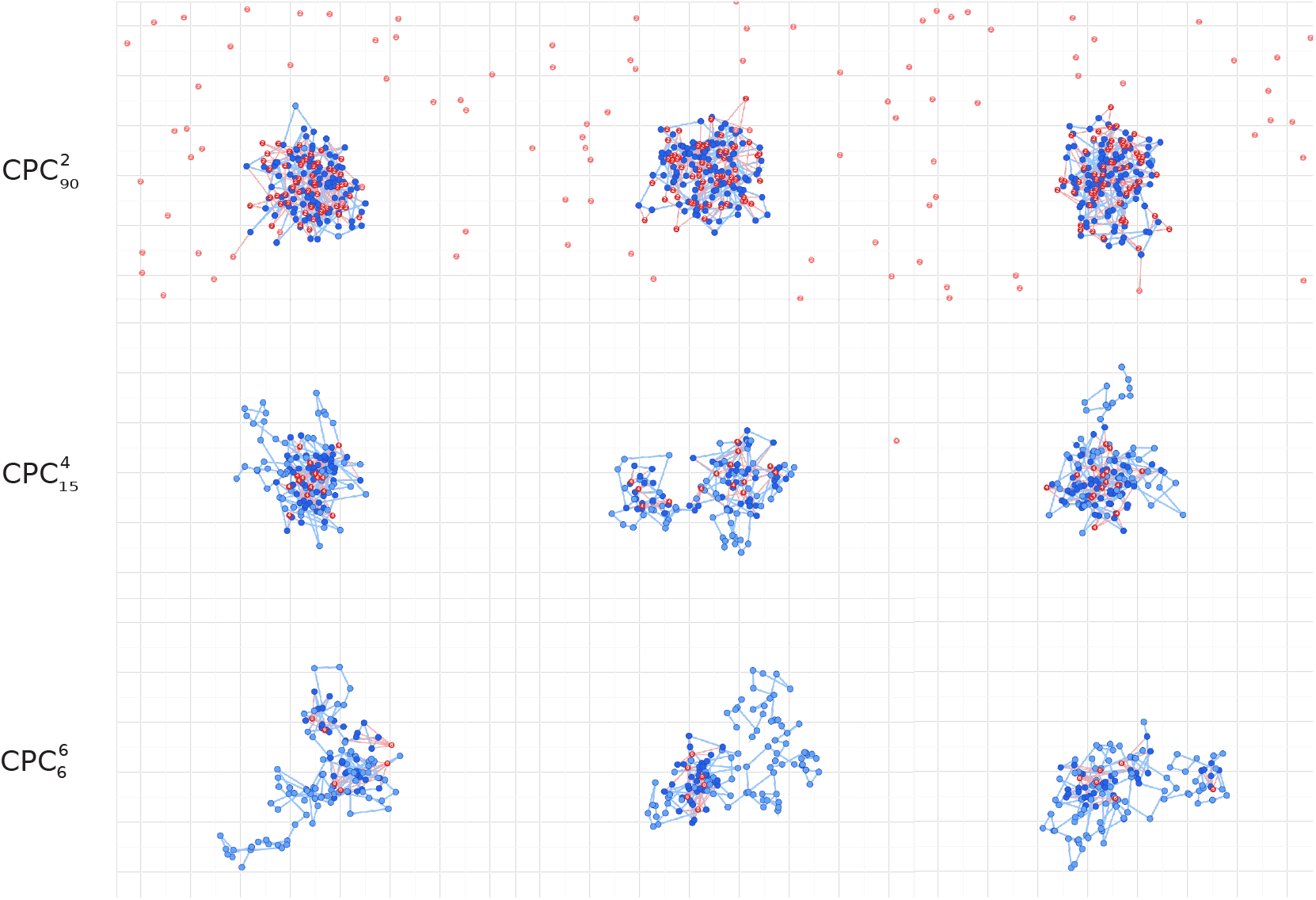
2D demonstrative model under a fixed connectivity budget *N*_*c*_ = 90. Representative steady-state chromatin conformations (blue beads) generated by the 2D Langevin-dynamics toy model with explicit CPCs of different valencies *k* (red beads). The total link capacity is held constant across *k* by adjusting CPC abundance, so differences in global shape primarily reflect the topology of constraints imposed by valency: low *k* yields more uniform compaction, whereas higher *k* promotes hub-like clustering and increased structural heterogeneity.

**Figure S12.**
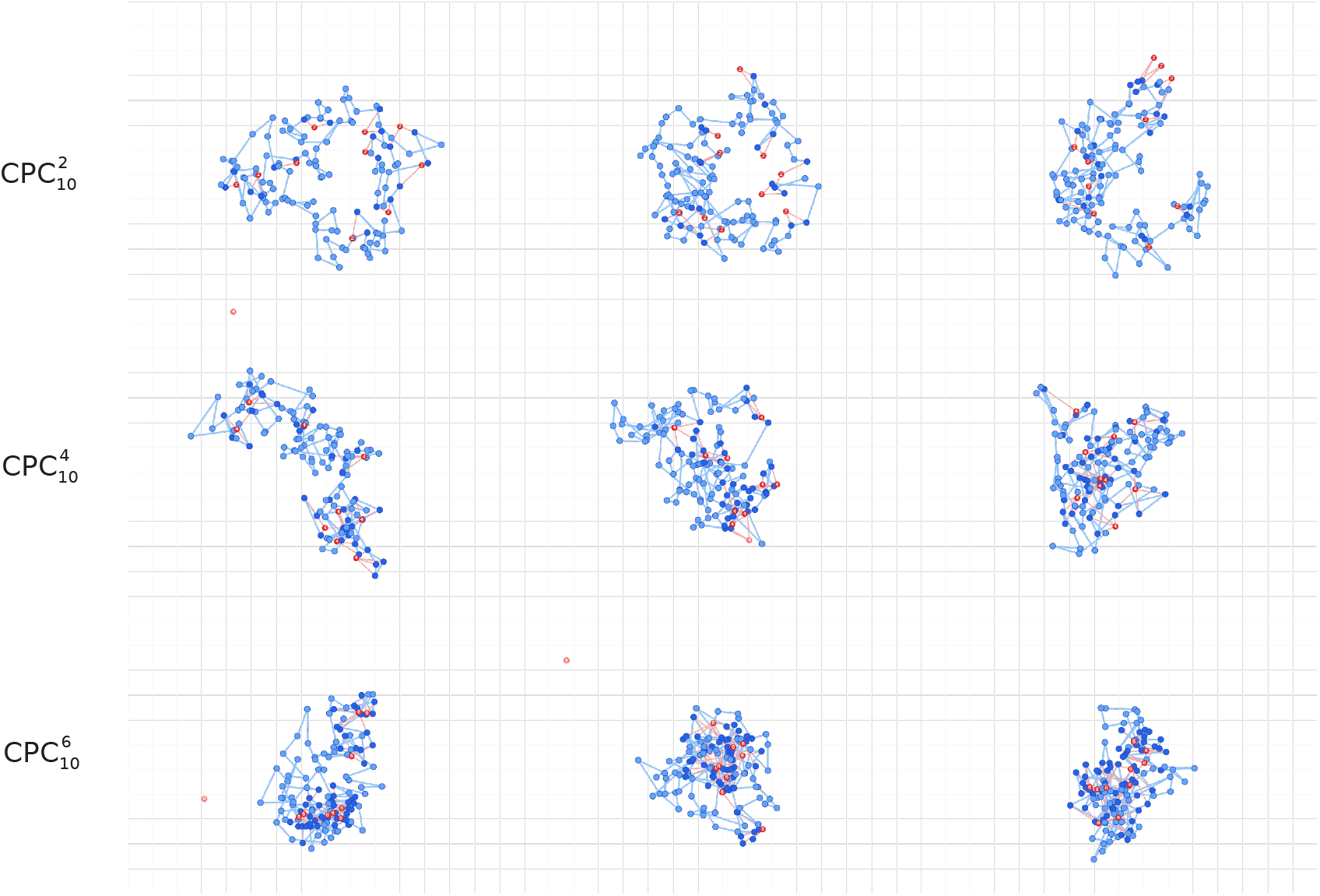
2D demonstrative model under a fixed CPC abundance *M* = 10 with the maximum link capacity increasing with valency, 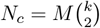. At low valency, the system is connectivity-limited and cannot sustain global compaction, whereas increasing *k* sharply increases the available links per CPC and drives stronger compaction with hub formation.

